# Self-iterative multiple instance learning enables the prediction of CD4^+^ T cell immunogenic epitopes

**DOI:** 10.1101/2025.02.02.636141

**Authors:** Long-Chen Shen, Yumeng Zhang, Zhikang Wang, Dene R. Littler, Yan Liu, Jinhui Tang, Jamie Rossjohn, Dong-Jun Yu, Jiangning Song

**Affiliations:** School of Computer Science and Engineering, Nanjing University of Science and Technology, 200 Xiaolingwei, Nanjing, 210094, China; Monash Biomedicine Discovery Institute and Department of Biochemistry and Molecular Biology, Monash University, Melbourne, VIC 3800, Australia; Monash Data Futures Institute, Monash University, Melbourne, VIC 3800, Australia; Department of Computer Science, Yangzhou University, Yangzhou, 225100, China; Institute of Infection and Immunity, Cardiff University, School of Medicine, Heath Park, Cardiff CF14 4XN, UK

**Author notes:** Correspondence should be addressed to: Jiangning Song,; Dong-Jun Yu;.

## Abstract

Accurately predicting the antigen presentation to CD4^+^ T cells and subsequent induction of immune response is fundamentally important for vaccine development, autoimmune disease treatments, and cancer neoepitope identification. In immunopeptidomics, single-allelic data are highly specific but limited in allele scope, while multi-allelic data contain broader coverage at the cost of weakly labeling. Existing computational approaches either overlook the massive multi-allelic data or introduce label ambiguity due to inadequate modeling strategies. Here, we introduce ImmuScope, a weakly supervised deep-learning framework integrating precise MHC-II antigen presentation, CD4^+^ T cell epitopes, and immunogenicity predictions. ImmuScope leverages self-iterative multiple-instance learning with positive-anchor triplet loss to explore peptide-MHC-II (pMHC-II) binding from weakly labeled multi-allelic data and single-allelic data, comprising over 600,000 ligands across 142 alleles. Moreover, ImmuScope can also interpret the MHC-II binding specificity and motif deconvolution of immunopeptidomics data. We successfully applied ImmuScope to discover melanoma neoantigens, revealing variations in pMHC-II binding and immunogenicity upon epitope mutations. We further employed ImmuScope to assess the effects of SARS-CoV-2 epitope mutations on immune escape, with its predictions aligned well with experimentally determined immune escape dynamics. Overall, ImmuScope provides a comprehensive solution for CD4^+^ T cell antigen recognition and immunogenicity assessment, with broad potential for advancing vaccine design and personalized immunotherapy.

## Introduction

T cell-mediated adaptive immunity is crucial for protection against pathogens and diseases^1–7^. Antigen presentation by major histocompatibility complex class II (MHC-II) molecules to CD4^+^ T cells is essential in initiating and coordinating a wide range of immune responses^8^. Experimental identification of CD4^+^ epitopes and characterization of MHC-II binding specificities are time-consuming and costly due to complex nature of antigen processing and the extensive polymorphism of MHC-II molecules^9^. Consequently, effective high-throughput prediction of CD4^+^ T cell epitopes, understanding MHC-II binding specificity, and assessing epitope immunogenicity are vital to developing vaccines and immunotherapies^10–15^.

Large-scale immunopeptidome datasets derived from liquid chromatography and mass spectrometry (LC-MS/MS)^16^ have significantly enhanced our understanding of MHC-II antigen presentation. Also known as eluted ligands (EL) datasets, they are categorized as single-allelic (SA) and multi-allelic (MA)^17, 18^ data, depending on whether allele-specific or pan-allelic antibodies are used during affinity purification. Single-allelic data provide precise, one-to-one peptide-MHC-II (pMHC-II) binding information. In contrast, multi-allelic data are weakly labeled, encompassing peptide interactions with multiple MHC-II alleles, where positive samples indicate binding to at least one allele, and negative samples indicate no binding. Multi-allelic data offer broader allele coverage, over twice as single-allelic data, especially for Human leukocyte antigens (HLA)-DQ and HLA-DP loci. Recent findings underscore the clinical relevance of previously underexplored molecules (e.g., HLA-DR3/4/5, HLA-DQ and HLA-DP) in autoimmune diseases^19–21^ and transplantation^22^, highlighting the necessity for integrated analysis of single- and multi-allelic data. Ultimately, incorporating weakly labeled multi-allelic data alongside single-allelic data mitigates SA-only biases and enables a comprehensive view of allele-specific binding patterns.

However, the weak labeling inherent in multi-allelic data (i.e., peptides are not directly assigned to specific allomorphs) presents unique challenges for model design and training. Most studies (e.g., HLAIImaster^23^ and BigMHC^24^) rely solely on single-allelic data, which restricts their ability to cover broader allele sets. Although Graph-pMHC^25^ and MixMHC2pred-2.0^26^ incorporate all possible pMHC-II pairs from multi-allelic data during training, they often generate numerous false positives. NNAlign_MA^27^ and NetMHCIIpan^17, 28^ leverage SA-trained neural networks to annotate multi-allelic data and are then fine-tuned with the pseudo-labeled data. However, prediction biases from the SA-based annotations can propagate through model training, particularly for alleles absent in single-allelic data. Furthermore, naive self-training strategies fail to capture the rich allelic diversity within multi-allelic data. Therefore, there is an urgent need to develop a highly precise model that effectively integrates both single- and multi-allelic data for predicting CD4^+^ T cell-related immunity.

Besides antigen presentation, numerous computational approaches have demonstrated impressive potential in predicting epitope characteristics and immunogenicity^18, 26, 29–33^. Nonetheless, the complexity of CD4^+^ T cell activation and differentiation still poses a hurdle^11, 34–36^. Most tools focus on a single facet of the cascaded immune process, such as NetMHCIIpan-4.0^17^ and NetMHCIIpan-4.2^30^ for antigen presentation, and DeepNeo^37, 38^ and TLimmuno2^39^ for epitope immunogenicity. HLAIImaster^23^, BigMHC^24^, and Graph-pMHC^25^ are designed to handle both facets, while MoDec^29^, NNAlign_MA^27^, MixMHC2pred-2.0^26^, and NetMHCIIpan-4.3^18^ can deconvolve MHC-II binding specificity. However, no studies have integrated the complete CD4^+^ T cell immune process—from antigen presentation and T cell recognition to immune response initiation within one framework^8^. Transferring cascaded immunological knowledge from the previous stage can enhance CD4^+^ T cell epitope predictions and help understand how the individual part of T cell immunity shapes the immune response^40–42^. Furthermore, current algorithms primarily lack fine-grained investigations across diverse immunopathological contexts or disease conditions, which may constrain their application potential in disease diagnosis and therapy^43, 44^.

Here, we propose ImmuScope, a weakly supervised deep-learning framework for CD4^+^ T cell immunity prediction, empowered by both single- and multi-allelic data. Utilizing the self-iterative multiple-instance learning approach and quality annotation filtering, ImmuScope pinpoints high-confidence pMHC-II pairs from weakly labeled multi-allelic data to broaden allele coverage. It utilizes a positive-anchor triplet loss to uncover more discriminative pMHC-II binding patterns. Trained on over 600,000 ligands spanning 142 MHC-II alleles, ImmuScope achieves state-of-the-art (SOTA) prediction accuracy with superior robustness and generalizability, expertly designed to navigate the complexities of highly polymorphic alleles in antigen presentation and immune response modeling. Moreover, ImmuScope serves as a comprehensive model that mirrors the cascade of T cell responses through transfer learning on multiple tasks, including antigen presentation prediction, MHC-II binding specificity discovery, CD4^+^ T cell epitope prediction, immunogenicity prediction, and motif deconvolution. We successfully apply ImmuScope to investigate the immunogenicity of melanoma neoantigens and assess the impact of epitope mutations on peptide-MHC-II binding. We further expand our analysis to SARS-CoV-2 epitopes, enabling the identification of key binding cores and exploration of immune escape mechanisms, particularly in the Omicron variant. Collectively, these findings underscore the broad applicability of ImmuScope in enhancing our understanding of T cell activation and advancing clinical applications in cancer and viral immunology.

## Results

### Overview of ImmuScope framework

We have developed ImmuScope, a weakly supervised deep-learning framework that integrates metric learning to robustly predict the CD4^+^ T cell responses. It supports a comprehensive suite of tasks including antigen presentation prediction, MHC-II binding specificity discovery, CD4^+^ T cell epitope prediction, immunogenicity prediction, and motif deconvolution, facilitating a thorough exploration of the cascaded immune process (**Fig. 1a**). Notably, the weakly labeled multi-allelic data spans a remarkably diverse range of allomorphs and contains massive peptide-binding data, particularly for the HLA-DQ and HLA-DP loci. Specifically, it comprises over 430,000 peptide samples—about 1.75 times that of the single-allelic data—and encompasses approximately 2.2 times as many MHC-II allomorphs (**Fig. 1b**). ImmuScope employs a multi-instance learning module to seamlessly integrate weakly labeled multi-allelic data with precisely annotated single-allelic data, thus harnessing the broad MHC coverage and extensive immunopeptidome of multi-allelic data alongside the specificity of single-allelic data. Furthermore, ImmuScope employs a metric learning loss function to capture more nuanced MHC-II binding specificities, thereby enhancing the discriminative power of the model.

**Fig. 1.**
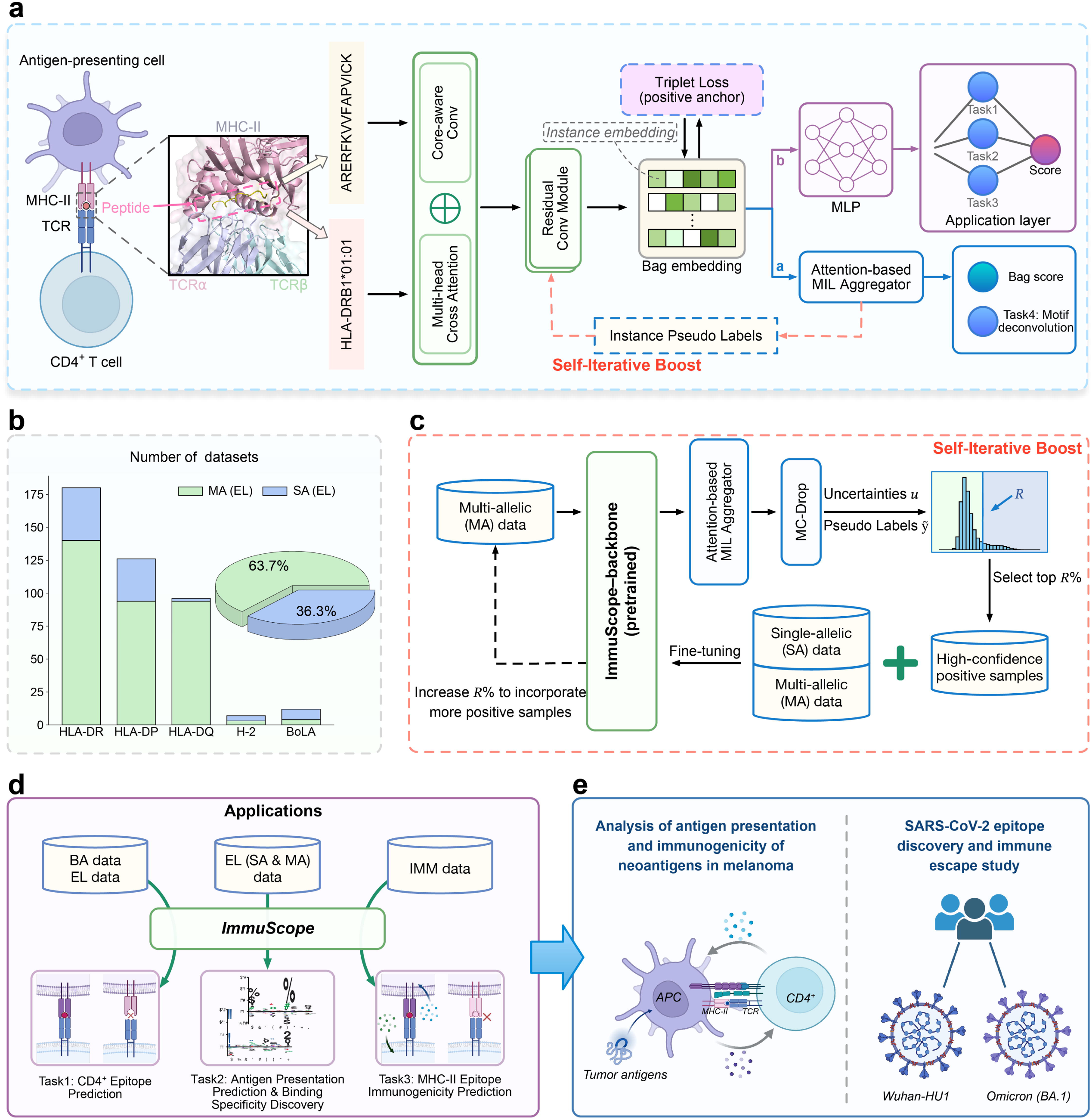
The overview of ImmuScope. **a**, Model backbone of ImmuScope consists of a pMHC-II interaction module (including core-aware convolution and multi-head cross-attention), a residual convolution module, and positive-anchor triplet loss. *branch a* focuses on learning bag-level features via an attention-based MIL aggregator, while *branch b* utilizes a multi-layer perceptron (MLP) to learn instance-level features for various application tasks. **b**, Numbers of multi-allelic (MA) and single-allelic (SA) data sets per MHC-II locus within the eluted ligand (EL) data, and the ratio of peptide counts for multi- and single-allelic data. **c**, Illustration of the self-iterative boosting strategy to generate high-quality pseudo labels for multi-allelic data and refine the model. **d**, Downstream tasks of ImmuScope and corresponding data inputs. **e**, Applying ImmuScope in cohort studies on melanoma neoantigens and SARS-CoV-2. Panels **a**, **d** and **e** created with <u>BioRender.com</u>.

The workflow of ImmuScope is as follows: Paired pMHC-II sequences are processed using a core-aware convolutional module and a multi-head cross-attention module to extract interaction embeddings (**Supplementary Fig. 1**). Residual convolutional blocks are used for further feature extraction to produce the final pMHC-II representations. To distinguish positive pairs from negative samples effectively, we employ a positive-anchor triplet loss, which is designed to maximize the distances of negative pairs and minimize the distances of positive pairs from an anchor, thereby enhancing model capacity without increasing computational burden. *Branch a* employs an attention-based MIL aggregator to learn individual allelic contributions within the multi-allele samples and compute the bag score. Meanwhile, *branch b* utilizes a multi-layer perceptron (MLP) adapted to the specific type of training data for predicting CD4^+^ T cell epitopes, antigen presentation, and immunogenicity. We further employ a self-iterative boosting strategy to select high-quality positive pseudo-labeled samples, which are then combined with single-allelic data to train the antigen presentation model, ImmuScope-EL (**Fig. 1c**). Recognizing that MHC-II-mediated antigen presentation is essential for initiating CD4^+^ T cell activation, we fine-tune ImmuScope-EL for other downstream tasks, including T cell epitope recognition and immunogenicity assessment (**Fig. 1d**). Furthermore, to assess model interpretability and practicality, we utilize ImmuScope to analyze antigen presentation and immunogenicity of neoantigens in a melanoma cohort and investigate SARS-CoV-2 epitope discovery and immune escape mechanisms (**Fig. 1e**).

### ImmuScope achieves state-of-the-art performance on CD4^+^ epitope benchmark

We evaluated the performance of ImmuScope and other algorithms, including Graph-pMHC^25^, MixMHC2pred-2.0^26^, NetMHCIIpan-4.2^30^, and NetMHCIIpan-4.3^18^, for identifying CD4^+^ T cell epitopes on the epitope benchmark. We employed them to predict the binding probability of each peptide to its given MHC-II allomorph and calculated the AUC for each source protein, epitope, and MHC-II allele entry. The AUCs demonstrated that ImmuScope significantly outperformed the current leading methods, NetMHCIIpan-4.3 and MixMHC2pred-2.0 (average AUC, 0.825 versus 0.771 and 0.761, respectively) (**Fig. 2a**). We also compared the average AUCs of ImmuScope and other methods across various HLA loci (**Extended Data Fig. 1a, b**), where ImmuScope exhibited better stability and superior performance. Additionally, we assessed the false positive rates using the Frank value, where ImmuScope showed enhanced performance in minimizing false positives, achieving a lower average Frank value of 0.181 (95% confidence interval [CI]: 0.166-0.195) compared to NetMHCIIpan-4.3 (0.229, 95% CI: 0.212-0.246) (**Fig. 2b** and **Extended Data Fig. 1c, d**). Further pairwise comparison with NetMHCIIpan-4.3 revealed that ImmuScope had a leading advantage in AUCs for 73.7% of the alleles and outperformed in Frank values for 61.4% of the alleles (**Fig. 2c**). We also analyzed the AUCs and Frank values of different methods across various peptide lengths (**Extended Data Fig. 1e, f**). Except for peptides of 16 amino acid length, where ImmuScope and NetMHCIIpan-4.3 exhibited comparable accuracy, ImmuScope revealed superior predictive performance at other peptide lengths. Overall, the benchmarking results indicate that ImmuScope exhibited an excellent capability of predicting CD4^+^ T cell epitopes.

**Fig. 2.**
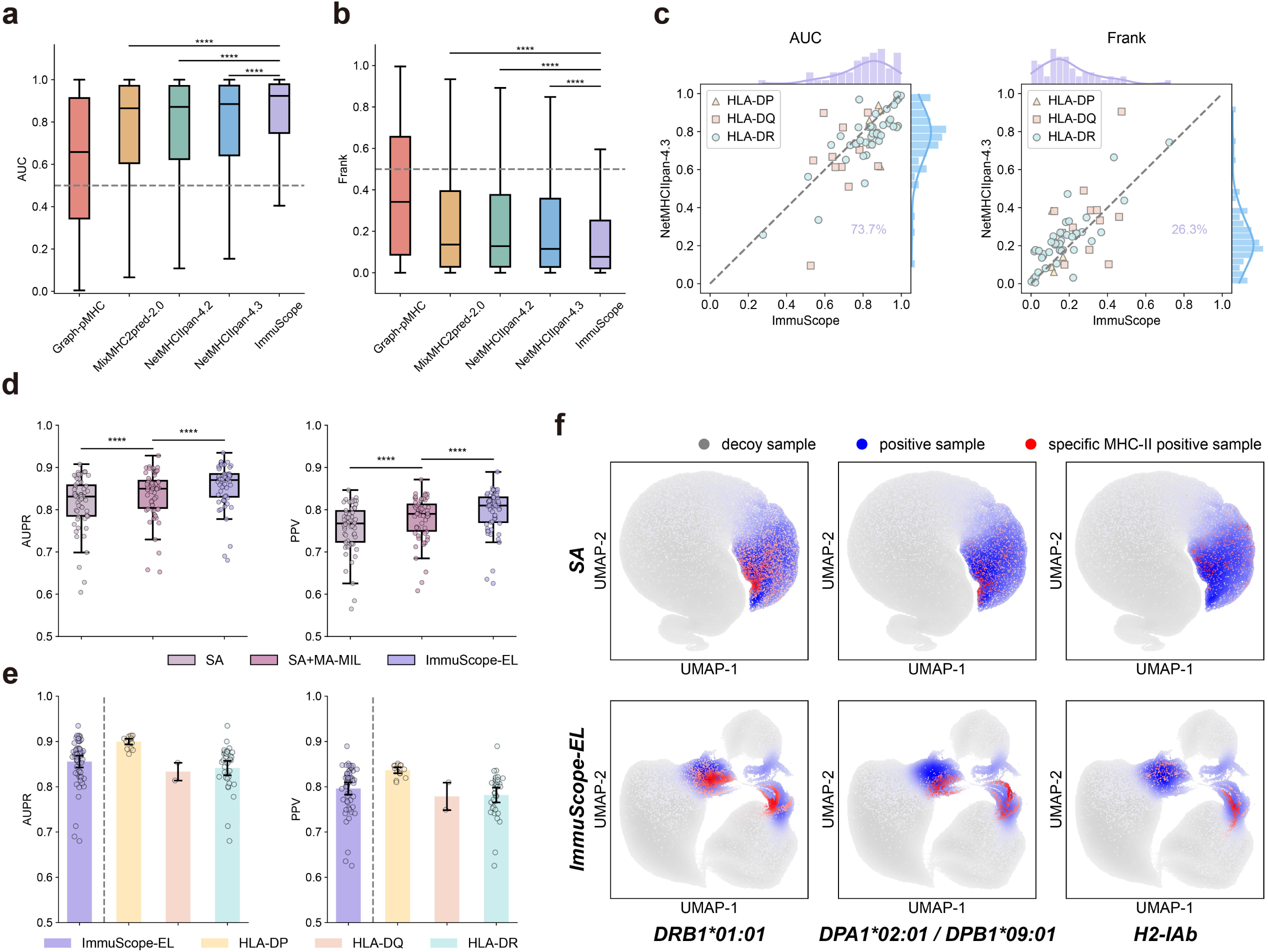
ImmuScope improves prediction of CD4^+^ T cell epitope and antigen presentation. **a**, Boxplot of AUCs on the CD4^+^ epitope benchmark. The *P* values were calculated by the paired Wilcoxon signed rank test to compare ImmuScope with existing methods (NetMHCIIpan-4.3, *P* = 1.9 x 10^-19^ ; NetMHCIIpan-4.2, *P* = 2.2 x 10^-19^ ; MixMHC2pred-2.0, *P* = 3.3 x 10^-22^, n=824). Box center line, median; box limits, upper and lower quartiles; whiskers, 1.5×interquartile range; dashed line, random; ****P < 0.0001. **b,** Boxplot of Frank values on the CD4^+^ epitope benchmark (NetMHCIIpan-4.3, *P* = 1.9 x 10^-19^ ; NetMHCIIpan-4.2, *P*= 2.3 x 10^-19^ ; MixMHC2pred-2.0, *P* = 4.4 x 10^-22^, n=824). **c**, Performance comparison between NetMHCIIpan-4.3 and ImmuScope based on (left) AUCs and (right) Frank values. **d**, Performance comparison across the SA, SA+MA-MIL, and ImmuScope-EL models: (left) AUPR and (right) PPV. ****P < 0.0001. **e**, AUPR and PPV of ImmuScope-EL by stratifying different HLA loci. The bars represent the mean by 1,000 bootstrap iterations, and the error bars indicate the 95% CIs. **f**, UMAP visualization of instance embeddings from the SA and ImmuScope-EL models on the test set. In **d**, **e**, each data point represents the performance of the corresponding HLA loci.

### Triplet loss and high-confidence pseudo-labels boost antigen presentation prediction

To evaluate the effectiveness of triplet loss and high-confidence pseudo-labels in enhancing antigen presentation prediction, we established three ablation experiments using 5-fold cross-validation (5cv): (1) an SA model was trained exclusively on single-allelic data through *branch b* and served as a baseline; (2) an SA+MA-MIL model incorporated triplet loss and MIL with single- and multi-allelic data but did not include iterative refining via positive pseudo-labels; (3) ImmuScope-EL additionally leveraged the high-quality positive pseudo-labeled data for model refining.

Integrating weakly labeled multi-allelic data through MIL substantially improved prediction accuracy while using triplet loss as an auxiliary function refined feature boundaries in the antigen presentation task, demonstrated by comparing the SA+MA-MIL model with the SA model across AUPR, PPV, and AUC0.1 (**Fig. 2d** and **Extended Data Fig. 2a**). Additionally, high-confidence positive pseudo-labels remarkably boosted model performance by comparing ImmuScope-EL and the SA+MA-MIL model (average AUPR, 0.856 versus 0.836; average PPV, 0.796 versus 0.779; average AUC0.1, 0.823 versus 0.805). Pairwise analysis of MHC-II alleles revealed that the SA+MA-MIL model consistently outperformed the SA model across almost all alleles (**Extended Data Fig. 2b and Supplementary Fig. 2**). Similarly, ImmuScope-EL showed explicit improvements over the SA+MA-MIL model. Moreover, we also illustrated the robustness of the ImmuScope-EL model using various indicators and stratifying different HLA loci (**Fig. 2e** and **Extended Data Fig. 2c**).

Next, we visualized the pMHC-II embeddings derived from the SA model and ImmuScope-EL model for multiple alleles, such as HLA-DRB1*01:01, HLA-DPA1*02:01/DPB1*09:01, H2-IAb, BoLA-DRB3*010:01 and BoLA-DRB3*020:02, by the UMAP^45^ algorithm (**Fig. 2f** and **Extended Data Fig. 2d**). Regarding the instances embeddings by ImmuScope-EL, both blue and red points representing positive samples are clustered in the manifold. The phenomenon of aggregation within the positive samples and separation between positive and negative samples is closely related to the positive-anchor triplet loss. Only positive samples were used as anchors since negative samples consisted of random natural peptides^18^ and was not a focus of our analysis. Consequently, introducing the auxiliary loss not only facilitates the formation of a bottleneck-like transition zone between positive and negative samples but also substantially enhances the discriminative capacity of the learned pMHC representations, thereby laying a stronger foundation for high-precision epitope identification and allele binding specificity analysis.

### Motif deconvolution on multi-allelic data with attention-based MIL module

We input the multi-allelic data into backbone of ImmuScope and then use the attention output of the MIL aggregator module and the antigen presentation score together for motif deconvolution analysis (**Fig. 3a**). Firstly, we performed 5cv experiments on the simulated multi-allelic dataset to assess the motif deconvolution efficacy of attention-based MIL due to the lack of precise labels on experimental multi-allelic data. The ensemble results for each MHC alleles achieved an average AUPR of 0.884 (95% CI: 0.882-0.885), an average AUC0.1 of 0.815 (95% CI: 0.814-0.817), and an average PPV of 0.819 (95% CI: 0.818-0.820) (**Fig. 3b**). To visually represent the impact of attention-based MIL deconvolution, we analyzed the corresponding clustering patterns of predicted attention scores and actual labels, indicating that ImmuScope successfully identified positive instances with high accuracy under weakly-supervised conditions (**Fig. 3c**). Consequently, leveraging high-confidence positive samples from multi-allelic data via motif deconvolution analysis could potentially refine antigen presentation prediction by broadening the available training information and increasing allele coverage.

**Fig. 3.**
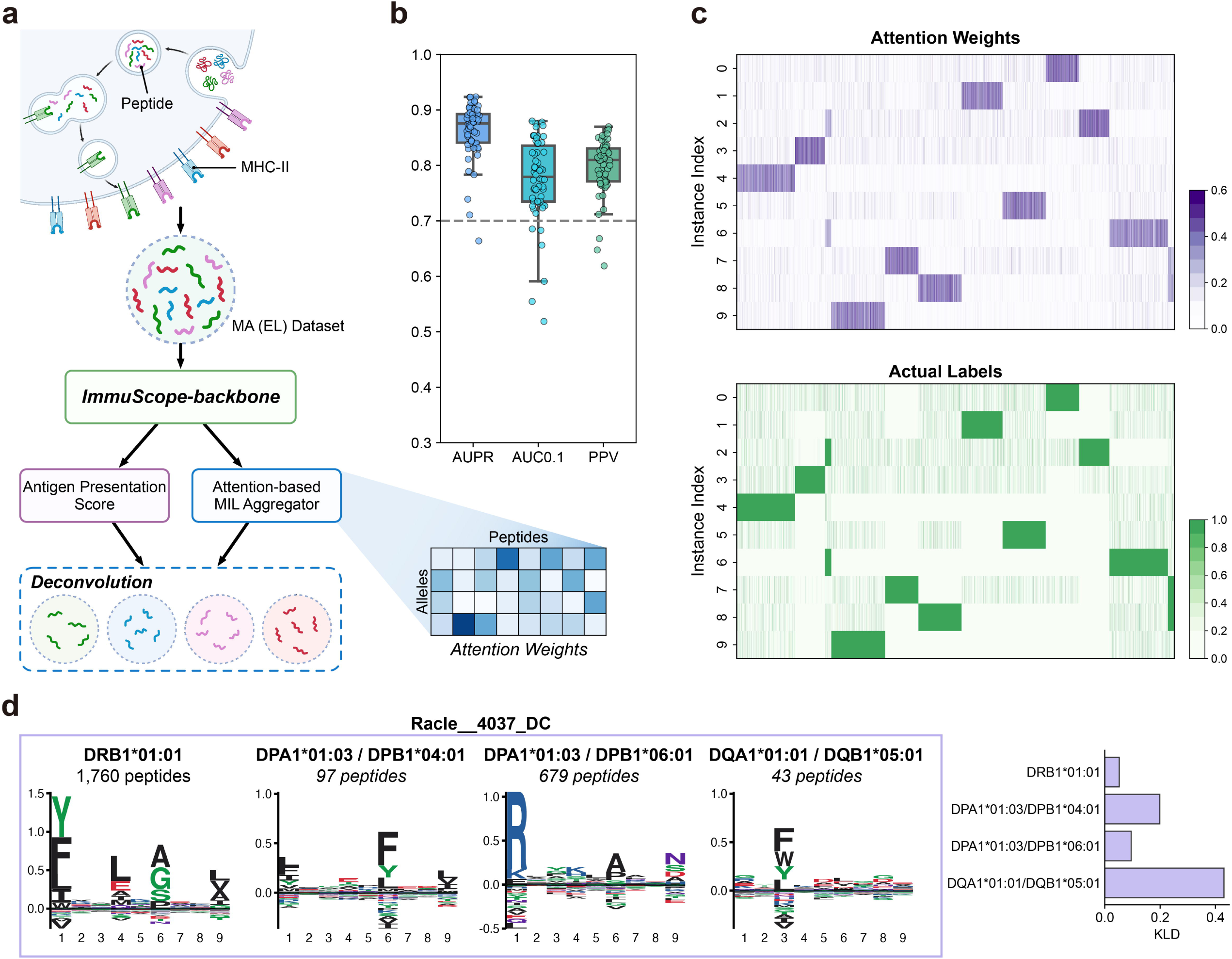
Motif deconvolution via ImmuScope on simulated and experimental multi-allelic data. **a**, Schematic diagram of MHC-II motif deconvolution pipeline. **b**, Performance (AUPR, AUC0.1 and PPV) of the attention-based MIL module across different MHC-II alleles in the simulated data. **c**, Heatmap of MIL attention weights and actual labels of the simulated data. **d**, Motif deconvolution logos on Racle 4037_DC heterozygous dataset and KLD analysis of PSFMs from the deconvoluted peptides and MHC-II immunopeptidomics. Panel **a** created with <u>BioRender.com</u>.

Next, we employed the attention-based MIL module on several subsets of the multi-allelic heterozygous dataset, which contain varying numbers of allele types, to identify potential alleles and represent their binding motifs via unsupervised alignment. Specifically, subsets Racle 4037_DC^29^, Racle RA957^29^, and Racle 3830_NJF_DQP^29^ include 4, 9, and 12 HLA alleles, respectively. Then, we use Seq2Logo^46^ to present the motifs derived from peptide sequences that bind to different MHC-II allomorphs recognized by our model in these heterozygous subsets. We particularly focused on HLA-DQ and HLA-DP molecules, which present fewer peptide ligands in the single-allelic data (**Fig. 3d** and **Extended Data Fig. 3a-c**). Comparison of these motifs with those cataloged in the MHC Motif Atlas (http://mhcmotifatlas.org/)^47^ revealed a high degree of similarity at the most conserved positions (**Supplementary Fig. 3a**). We computed the Kullback-Leibler divergence (KLD) between the position-specific frequency matrices (PSFMs) generated from the deconvoluted peptides and those derived from immunopeptidomics datasets. The binding motifs across most of the alleles except HLA-DQA1*01:01/DQB1*05:01 demonstrated remarkably high similarity to Racle 4037_DC heterozygous datasets (**Fig. 3d**). Limited peptide ligands might result in less clearly defined motifs, such as for HLA-DQA1*01:01/DQB1*05:01. Additionally, ImmuScope-EL successfully inferred motifs for MHC-II alleles absent from the database comparable to those found using NetMHCIIpan-4.3, e.g. HLA-DQA1*03:03/DQB1*04:02 and HLA-DQA1*05:05/DQB1*03:02 (**Extended Data Fig. 3a, b and Supplementary Fig. 3b**). These findings underscore ImmuScope-EL’s utility as a computational tool for identifying binding peptides within multi-allelic data and analyzing binding motifs of the MHC-II alleles.

### ImmuScope quantifies MHC-II binding specificities for allomorphs without known ligands

MHC-II polymorphism might impede the prediction of pan-allelic binding specificity. To assess the performance of ImmuScope-EL on unseen alleles, we compared it with current state-of-the-art methods, such as NetMHCIIpan-4.3 and MixMHC2pred, in predicting allelic binding specificity across different loci. We employed a leave-one-allele-out cross-validation strategy, i.e., excluding the alleles under prediction from the training dataset. Subsequently, we generated PSFMs for each allele through predicting 100,000 random human peptides by ImmuScope-EL and selecting the top 1% with the highest binding scores (**Fig. 4a**). We then evaluated the consistency (using KLD distance) between the PSFMs predicted by ImmuScope-EL or other predictors and those sourced from the MHC-II immunopeptidomics studies. ImmuScope-EL generally demonstrated superior capability in inferring the binding specificity of allomorphs lacking known peptide ligands (**Fig. 4b**). Notably, ImmuScope-EL enhanced predictive accuracy for the multiple specificities of HLA-DRB1*08:02 and the bidirectional-binding specificity^48^ of HLA-DPA1*02:01/DPB1*09:01.

**Fig. 4.**
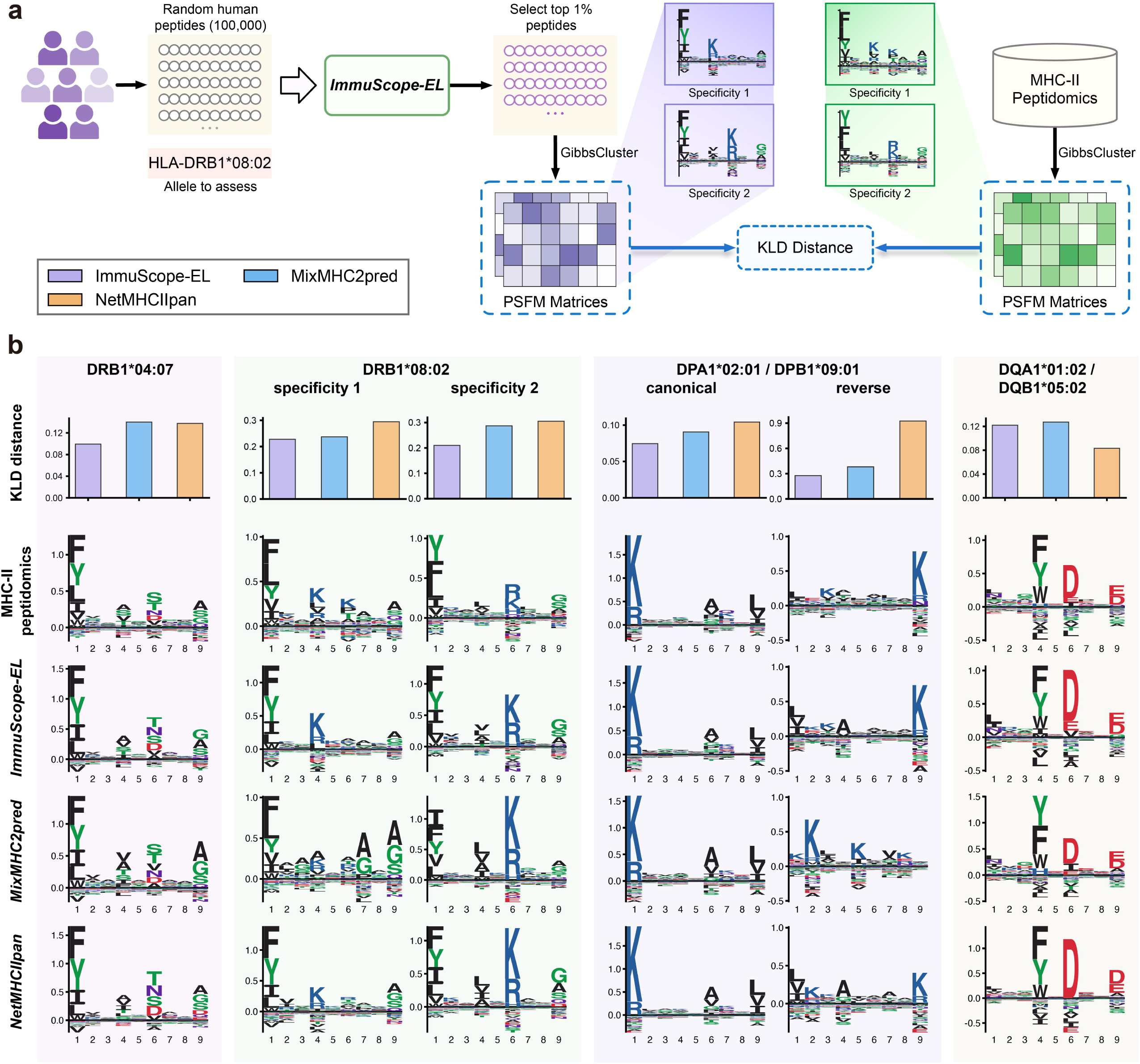
MHC-II binding specificities for allomorphs without known ligands predicted by ImmuScope. **a**. The workflow of MHC-II binding specificity evaluation. **b**, Comparison of ImmuScope with other algorithms in predicting MHC-II binding specificity. Bar plots of KLD distance against the PSFMs from the MHC-II immunopeptidomics data (top); peptide binding motifs obtained by different methods (bottom). Motif of HLA-DQA1*01:02/DQB1*05:02 are derived from peptide ligands in the MHC Motif Atlas Database (http://mhcmotifatlas.org/). Other MHC-II peptide ligands were derived from the corresponding positive samples in the single-allelic data. Panel **a** created with <u>BioRender.com</u>.

Despite the constraint of a limited dataset, which included only two single-allelic HLA-DQ sets, ImmuScope-EL successfully predicted the binding specificity of HLA-DQA1*01:02/DQB1*05:02. The prediction was corroborated by data from the MHC Motif Atlas database, reinforcing the reliability of our approach. The binding motifs highlighted the substantial disparities in sequence binding propensities between HLA-DQA1*01:02/DQB1*05:02 and HLA-DQ allomorphs in the training set, e.g., HLA-DQA1*01:02/DQB1*06:02 and HLA-DQA1*01:02/DQB1*06:04 (**Supplementary Fig. 4**). These findings illustrate ImmuScope-EL’s capability to elucidate local binding patterns across alleles, adeptly addressing the extensive polymorphism characteristic of MHC-II alleles.

### ImmuScope enhances epitope immunogenicity prediction accuracy

Accurate prediction of immunogenic peptides that activate CD4^+^ T cells is essential for both vaccine development^49^ and immunotherapy^50^. To demonstrate the effectiveness of ImmuScope-IM in predicting immunogenicity, we compared it with five existing algorithms, including DeepNeo^37^, MixMHC2pred-2.0^26^, NetMHCIIpan-4.3^18^, TLimmuno2^39^, and HLAIImaster^23^. Notably, ImmuScope-IM exhibited superior performance on the immunogenicity benchmark with an overall AUC of 0.909 (95% CI: 0.901-0.918) (**Fig. 5a**). We further assessed the predictions across different MHC-II alleles. For MHC-II alleles with a sample size greater than 10 and at least one immunogenic epitope, ImmuScope-IM consistently showed statistically significant improvements of AUC over TLimmuno2 and HLAIImaster with *P* values of 1.4 x 10^-7^ and 2.2 x 10^-7^, respectively (**Fig. 5b**). ImmuScope-IM outperformed HLAIImaster in 89.1% of the MHC-II alleles (**Fig. 5c**). Given the prevalence of fewer positive samples in real-world scenarios, we adopted a positive to negative ratio of 1:10 in building the immunogenicity dataset. Considering the imbalance of data, the precision-recall curve (PRC) was utilized for a more accurate assessment of the model performance. The AUPRs highlighted significant improvements of ImmuScope-IM over existing methods (**Extended Data Fig. 4a, b**). Predicted AUPRs for ImmuScope-IM and the leading existing model, HLAIImaster, were compared across different MHC-II alleles (**Extended Data Fig. 4c**). ImmuScope-IM demonstrated superior or comparable performance compared to HLAIImaster in 92.7% of the 55 MHC-II alleles.

**Fig. 5.**
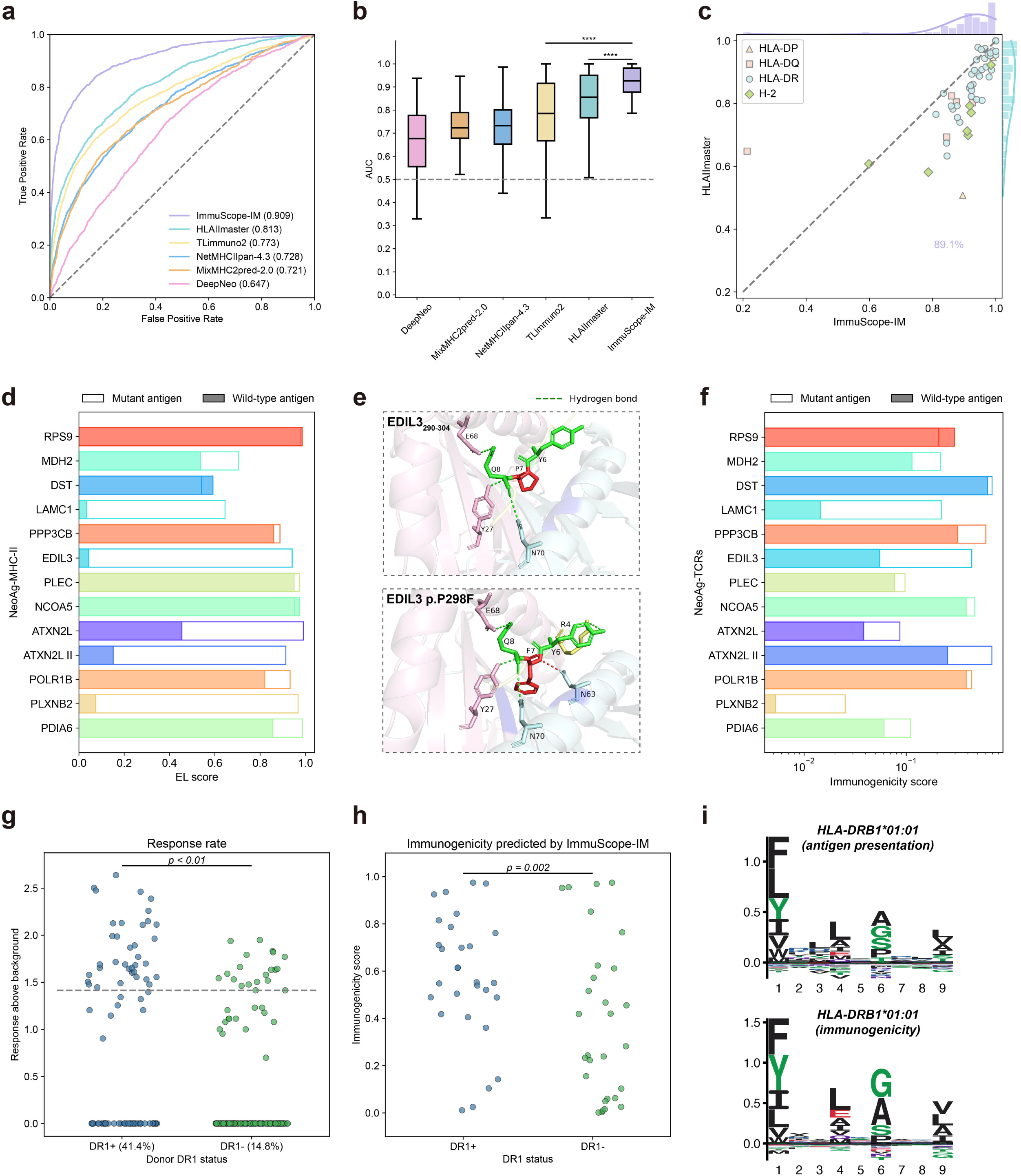
Immunogenicity benchmarking and analysis of melanoma neoantigens and SARS-CoV-2 epitopes. **a**, ROC curves of ImmuScope-IM and other methods on the immunogenicity benchmark. **b**, AUCs of ImmuScope-IM and other methods on the immunogenicity benchmark. The *P* values were calculated by the paired Wilcoxon signed rank test to compare ImmuScope-IM with existing methods (HLAIImaster, *P* = 2.2 x 10^-7^ ; TLimmuno2, *P* = 1.4 x 10^-7^, n=62). Box center line, median; box limits, upper and lower quartiles; whiskers, 1.5× interquartile range; points, data points; ****P < 0.0001 **c**, Pairwise comparison of AUCs between HLAIImaster and ImmuScope across different MHC-II alleles. **d**, Predictive analysis of melanoma neoantigen presentation based on ImmuScope-EL. **e**, Structural conformation of EDIL3_290-304_ epitopes bound to HLA-DPA1*01:03/DPB1*02:01 upon the mutation predicted by AlphaFold3 (average pLDDT=92.5, ipTM=0.92). The amino acid at the mutation site is marked in red. Dashed lines indicate the hydrogen bonds, and interaction sites within 4 Å are displayed in dark blue. **f**, Predictive analysis of the immunogenicity of melanoma neoantigens based on ImmuScope-IM. **g**, Summary of maximal responses from all peptides (total n=29) across all donors tested (total n=8), categorized by donor DR1 status, with responses scaled on y-axis as 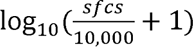. Each point represents a donor’s peak response to a peptide from two ELISpot wells. The dashed line at 1.415 notes the cutoff for donor-peptide responses, representing 25 sfcs/10,000 cells. The response rate is expressed as a percentage. An inset shows the P-value was calculated using Fisher’s exact test, to compare the DR1^+^/DR4^+^ statuses (groups) with the positive peptide responses. **h**, Summary of DR1^+^/DR4^+^ status predictions by ImmuScope-IM for all peptides (n=29). The *P* values was calculated using paired Wilcoxon signed rank test to compare the peptide responses associated with DR1^+^/DR4^+^ status. **i**, Predicted binding peptide motifs for HLA-DRB1*01:01 as determined by the antigen presentation (ImmuScope-EL) and immunogenicity (ImmuScope-IM) models, respectively.

### ImmuScope precisely reveals immunogenic neoantigens in melanoma

To investigate the practicality of ImmuScope for neoantigen identification, we applied it to a cancer cohort of cutaneous melanoma^51^ (**Supplementary Table 1, 2**). ImmuScope effectively detected the HLA class II-presented and immunogenic neoantigens within the tumor microenvironment and facilitated the evaluation of clinical outcomes. Initially, ImmuScope-EL was utilized to predict the likelihood of various HLA class II allomorphs in Pt-C and Pt-D binding to specific neoantigens, and to identify the most likely HLA class II restrictions and binding cores (**Extended Data Fig. 4d, e and Supplementary Fig. 5**). Our predictions aligned closely with those reported by Oliveira et al.^52^ using NetMHCIIpan 4.0 except for the antigen EDIL3. HLA-DPA1*01:03/DPB1*02:01 and HLA-DPA1*01:03/DPB1*04:02 were both predicted to present EDIL3 with high probability. These allomorphs, differing by merely two amino acids, exhibit considerable sequence homology and robust antigen presentation abilities. Further analysis revealed distinct differences in antigen presentation between mutant and wild-type antigens, particularly concerning their structural organizations around the mutation site (**Fig. 5d**). Notably, neoantigens that underwent mutations at key MHC-II anchor positions exhibited substantial changes in antigen presentation probability compared to wild-type antigens, as seen in LAMC1, EDIL3, ATXN2L II, and PLXNB2. The wild-type epitopes failed to elicit an immune response partly due to inadequate biophysical conditions for peptide-MHC-II binding.

Moreover, we predicted the interaction conformations between EDIL3_290-304_ and HLA-DPA1*01:03/DPB1*02:01 pre- and post-mutation by AlphaFold3 (**Fig. 5e, Supplementary Table 3**). The substitution of proline with phenylalanine significantly enhanced the peptide’s binding affinity to MHC-II, primarily due to the increased hydrophobicity and bulkier size, which improved its stabilization in the MHC binding groove. Additionally, the main chain of phenylalanine formed an extra hydrogen bond, further stabilizing the peptide-MHC interaction and enhancing the peptide’s conformational adaptability. The structural evidence revealed a better fit with the groove’s shape and chemical properties of MHC, likely boosting the peptide’s presentation efficiency. Using ImmuScope-IM, we also compared the T cell activation levels between mutant and wild-type antigens (**Fig. 5f**). Most of the neoantigens demonstrated enhanced immunogenicity compared to their wild-type counterparts expect for RPS9. These results corroborate the high concordance of ImmuScope-predicted neoantigen characteristics with experimental assays regarding antigen presentation, binding core, and immunogenicity, underscoring the potential of ImmuScope for advancing tumor neoantigen discovery.

### ImmuScope predictions aligns closely with SARS-CoV-2 epitope discovery results

We further assessed the application potential of ImmuScope-IM in clinical cohorts via a longitudinal study ^53^. Chen et al. identified and characterized CD4^+^ T cell epitopes derived from SARS-CoV-2 and restricted by the prevalent HLA-DR1 allotype. Their analysis illustrated the maximal immune responses of eight donors to 29 candidate SARS-CoV-2 epitopes (**Fig. 5g**). Immune responses were quantified as spot-forming cells (SFCs) per 10,000 PBMCs. The study initially utilized HLA-DR1 (DRB1*01:01) as the positive allotype for assessing peptide binding and immunogenicity, while HLA-DR4 (DRB1*04:01) served as the negative allotype. Blood samples from donors both with and without these specific HLA types were tested to evaluate the *in vitro* immunogenicity of the peptides. We utilized ImmuScope-IM to predict the immunogenicity probabilities for all peptides in the context of both DR1^+^ and DR4^+^ status (**Fig. 5h**). Additionally, we conducted a paired Wilcoxon signed rank test, yielding a P-value of 0.002. These findings are consistent with the experimental results reported by Chen et al., suggesting that HLA-DR1 donors exhibit enhanced immune responses.

We also predicted the antigen presentation and immunogenicity binding motifs of HLA-DRB1*01:01 (**Fig. 5i**). We accurately identified the peptide-binding groove of HLA-DR1 epitopes derived from SARS-CoV-2, as defined structurally by Chen et al., through computational alignment scores (**Fig. 6a**, **b**, and **Extended Data Fig. 5a-d**). Notably, we identified a more closely matching core binding site on the spike protein epitope S_486-505_. The complex structure of the binding peptide *LQSYGFQPTNGVGY* with HLA-DRB1*01:01 was predicted by AlphaFold3 with a predicted local distance difference test (pLDDT) score greater than 90 and an interface predicted TM score (ipTM) of 0.94 (**Supplementary Table 3**). The predicted protein complex structure exhibited exceptionally high local accuracy and interface alignment quality, rendering it highly reliable and instrumental in identifying potential immunogenic epitopes.

**Fig. 6.**
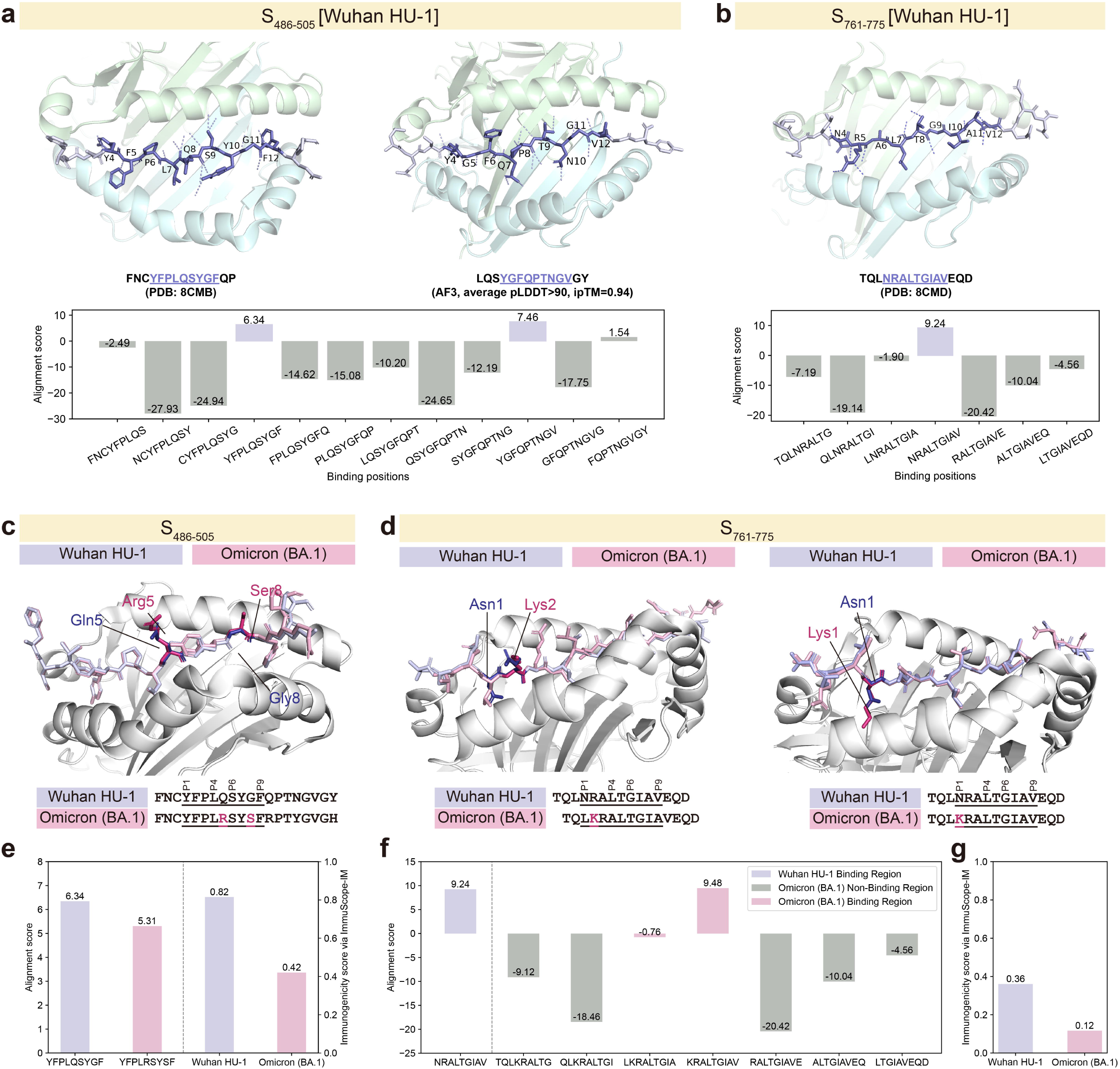
Predictive analysis of the spike epitope binding core and the structural variations of Omicron (BA.1). a,b, Predicted binding positions and alignment scores by ImmuScope-EL for SARS-CoV-2 spike protein epitope S_486-505_ (**a**) and S_761-775_ (**b**), respectively. The residues of the binding core within each peptide are labeled according to their positions. The interactions between the peptides and HLA-DR1 are displayed by dashed lines. **c**, Structural comparison of HLA-DR1-S_486-505_^Omicron^ ^(BA.1)^ aligned on the HLA-DR1-S_486-505_^Wuhan^ ^HU-1^ structure. The HLA-DR1 peptide-binding groove is depicted as gray cartoon, while the S_486-505_^Omicron^ ^(BA.1)^ and S_486-505_^Wuhan^ ^HU-1^ peptide are displayed as pink and lavender sticks. The mutant amino acids are also highlighted. **d**, Structural comparison of HLA-DR1-S_761-775_^Omicron^ ^(BA.1)^ aligned on the HLA-DR1-S_761-775_^Wuhan^ ^HU-1^ structure in two registers: (left) a+1 register shift in asymmetric unit (ASU) copies 1 and 3; (right) the same register as seen in ASU copy 2^53^. **e**, Alignment scores for the binding core of HLA-DR1-S_486-505_ ^Wuhan^ ^HU-1^ and ^Omicron^ ^(BA.1)^ predicted by ImmuScope-EL and the corresponding immunogenicity scores predicted by ImmuScope-IM. **f**, Alignment scores for the binding core of HLA-DR1-S_761-775_^Wuhan^ ^HU-1^ and various binding positions of Omicron (BA.1) predicted by ImmuScope-EL. **g**, Immunogenicity scores of HLA-DR1-S_761-775_^Wuhan^ ^HU-1^ and ^Omicron^ ^(BA.1)^ predicted by ImmuScope-IM.

### ImmuScope facilitates to understand SARS-CoV-2 immune escape dynamics

Furthermore, Chen et al. delineated the influence of SARS-CoV-2 variant mutations on epitope presentation and clarified immune evasion mechanisms by performing crystallographic analyses on HLA-peptide complexes of variant epitopes, revealing their strategies for eluding memory CD4^+^ T cell detection. The study showed that despite increased binding to HLA-DR1, the S_486–505_ and S_761–775_ peptides of Omicron (BA.1) were not recognized by T cells. We utilized ImmuScope-IM to investigate changes in the binding cores and immunogenicity of SARS-CoV-2 epitopes upon the mutations.

Structural analysis of the HLA-DR1-S_486-505_^Wuhan^ ^HU-1^ revealed that the S_486-505_^Omicron^ ^(BA.1)^ induced mutations located within both the binding core and the peptide flanking region (PFR). HLA-DR1 bound the S_486-505_^Omicron^ ^(BA.1)^ epitope using the same register (**Fig. 6c**), which was consistent with the core binding alignment scores predicted by ImmuScope-EL (**Fig. 6e**). The core sequence *YFPLRSYSF* exhibited a slight reduction in post-mutation binding score, indicating that the S_486-505_^Omicron^ ^(BA.1)^ maintained favorable binding affinity with HLA-DR1. Specifically, all core-positioned mutations occurred at potential TCR contact positions, notably at non-anchor residues Q493R (P5) and G496S (P8). Q493R (P5) introduced the most evident conformation changes, where P5-Arg with positive charges was located at the center of the binding core. G496S introduced an additional polar hydroxyl at position P8-Ser. The immunogenicity scores predicted by

ImmuScope-IM decreased from 0.815 to 0.420 after mutation, which was consistent with the immune escape mechanisms of S_486-505_^Omicron^ ^(BA.1)^ according to the structural analysis (**Fig. 6e**). In addition, we analyzed the case of register shift caused by a single mutation in the S_761-775_^Omicron^ ^(BA.1)^ by ImmuScope. N764K, positioned at archetypal P1 anchor position for HLA-DR1, resulted in two distinct peptide conformations of the HLA-DR1-S_761-775_^Omicron^ ^(BA.1)^ (**Fig. 6d**). In the first conformation, the HLA molecule contacted the neighboring Leu at the P1 pocket position and generated a+1 register frameshift (i.e., *TQL<u>KRALTGIAV</u>EQD* to *TQ<u>LKRALTGIA</u>VEQD*). This new peptide conformation bound to HLA-DR1 via P1-Leu, P4-Ala, and P9-Ala, and the unfavored Thr at P6 due to its large and polar hydroxyl side chain^53^. The second conformation accommodated the N764K mutation at P1 to bind S ^Omicron^ ^(BA.1)^, aligning with the binding register of S_761-775_^Wuhan HU-1^.

We also calculated the alignment scores of binding regions upon the mutation. *KRALTGIAV* achieved the highest score, corresponding to the second peptide conformation. The alignment score of *LKRALTGIA* was -0.758 (**Fig. 6f**) and 1.574 when excluding the P6 position (**Extended Data Fig. 5e**). *LKRALTGIA* still exhibited a suboptimal binding probability relative to other regions, which aligned with the structural analysis^53^. Compared to the pre-mutation immunogenicity score of HLA-DR1-S_761-775_^Wuhan^ ^HU-1^, the immunogenicity score of HLA-DR1-S ^Omicron^ ^(BA.1)^ predicted by ImmuScope-IM was reduced by 67.8% (**Fig. 6g**). This reduction might account for the structural and sequence alterations caused by the frameshift mutation, which further affected T cell recognition.

## Discussion

ImmuScope presents a comprehensive framework that seamlessly integrates weakly supervised learning with metric learning to enable the accurate prediction of CD4^+^ T cell-mediated immune responses, offering key insights for the development and evaluation of precision immunotherapies. Unlike previous algorithms, ImmuScope employs self-iterative multiple-instance learning to combine weakly labeled multi-allelic data and highly specific single-allelic data, achieving over a two-fold increase in allele coverage. It also incorporates positive-anchor triplet loss to effectively identify challenging pMHC-II samples to improve MHC-II binding specificity. This synergistic approach empowers ImmuScope to achieve SOTA performance in antigen presentation prediction, T cell epitope recognition, and epitope immunogenicity assessment, while also providing reliable MHC-II binding specificity analysis and motif deconvolution. Moreover, ImmuScope demonstrates great potential for assessing the immunogenicity of melanoma neoepitopes and their antigen presentation levels, as well as for revealing the mutation effects of SARS-CoV-2 epitopes on antigen presentation and immune escape. As high-quality data on antigen presentation and immunogenicity continue to accumulate, ImmuScope is expected to become a crucial tool for accelerating vaccine development and optimizing immunotherapeutic strategies.

Although ImmuScope outperforms other methods in mining weakly labeled multi-allelic data and interpretability, there is still room for improvement in real-world clinical applications, especially in predicting the immunogenicity of T cell epitopes. In this study, we exclusively used IFN-*γ* to determine the immunogenicity of CD4^+^ T cells to specific peptides, in accordance with mainstream algorithms, which introduces inherent limitations. Although IFN-*γ* is a crucial Th1-type cytokine essential for mediating cellular immune responses, it does not fully capture the multifunctionality of CD4^+^ T cells or the complexity of immune responses. For instance, it does not reveal other types of immune responses such as those regulated by cytokines like IL-4, IL-10, and IL-17, which are Th2 and Th17 responses^54, 55^. Additionally, relying solely on IFN-*γ* cytokine does not allow us to assess T cell proliferation or the expression of phenotypic markers such as CD25 and CD69, which are important indicators of T cell activation and function^55^. Although ImmuScope effectively utilizes large-scale pMHC-II sequence data to assess CD4^+^ T cell epitopes and immunogenicity, integrating the extremely limited structural data on TCR-pMHC-II or pMHC-II complexes remains challenging. Currently, the TCR3d^56^ database includes only about 300 experimentally determined structures of TCR-pMHC-II or pMHC-II complexes in both bound and unbound forms. The scarcity of structural data significantly restricts our ability to use advanced deep learning techniques to accurately model and infer the intricate interactions between TCR and pMHC-II, thereby limiting the possibility of obtaining direct molecular understanding of the immune recognition mechanism.

Future studies should focus on incorporating a broader range of immunological knowledge, including cytokine profiles, cell phenotypes, and proliferation data. Advances in high-resolution structure determination technology and the development of structure prediction algorithms like AlphaFold3 are expected to significantly increase the availability of structural information on TCR-pMHC-II complexes^57, 58^. In upcoming work, we aim to expand the ImmuScope model by seamlessly integrating sequence and structural data, thereby enabling a precise elucidation of the molecular interaction mechanisms of pMHC-II. This enhancement will facilitate the development of a more holistic and comprehensive T cell immune epitope prediction framework. These advancements will establish a robust scientific foundation, driving the innovation and efficacy of next-generation vaccines and immunotherapies^59, 60^.

## Methods

### Datasets

The statistics of the datasets used for training and validation for different tasks are shown in **Supplementary Table 4**. The following is a detailed dataset description:

#### MHC-II antigen presentation data

To train the antigen presentation model, we used the large-scale antigen presentation data collected by Nilsson et al.^17, 18, 30, 61^, comprising three data types: binding affinity (BA), single- and multi-allelic eluted ligand (EL) datasets (**Fig. 1b** and **Supplementary Fig. 6a, b**). All data were filtered to remove possible contaminants and MHC class I restricted peptides, retaining peptides of 12-21 amino acids (AAs) in length^26^. The EL datasets were then enriched by uniformly sampling five times of 12-21 AAs random natural peptides as negative samples. The datasets were divided into five subsets for cross-validation (CV) using the common-motif method, ensuring that peptides sharing a subsequence of nine or more amino acids were grouped into the same subset^62^. The final single-allelic dataset contains 246,590 positive and 2,448,316 negative samples, while the multi-allelic dataset includes 432,255 positive and 4,467,755 negative samples, covering a total of 142 MHC-II molecules. Additionally, the binding affinity dataset comprises 129,110 data points across 80 class II molecules.

#### CD4^+^ epitope benchmark

The CD4^+^ epitope benchmark^18^, compiled by Nielsen et al. in 2023, was assembled following a specific protocol. Initially, positive CD4^+^ T cell epitopes ranging from 12-21 amino acids, without post-translational modifications and with complete 4-digit MHC-II typing, were selected from the Immune Epitope Database (IEDB, https://www.iedb.org/)^63^. Only epitopes associated with well-documented source proteins were considered. Subsequently, corresponding negative samples were generated based on the source protein sequences retrieved from the UniProt database (https://www.uniprot.org/)^64^. Each {epitope, allele, protein} triplet was then segregated into a distinct test subset. Within each subset, using a sliding window of the same length as the epitope, overlapping peptides were generated from the source protein sequence and designated as negative samples, excluding the epitope itself. Furthermore, it was ensured that none of the samples in the test set had previously appeared in the MHC-II antigen presentation training data. Ultimately, the test set comprised 842 {epitope, allele, protein} triplets, encompassing 40 HLA-DR, 13 HLA-DQ, and 4 HLA-DP molecules.

#### Immunogenicity data

We curated immunogenicity assay data from IEDB^63^ and integrated it with the MHCBN^65^ dataset, following the methodology described in DeepNeo^37, 38^. This dataset contains records up to May 14, 2024. Specifically, we selected data of T cell reactivity based on IFN-*γ* secretion. Furthermore, we refined the dataset to include entries only with full MHC-II restriction and peptide lengths ranging from 12 to 21 AAs. Given the variable nature of pMHC-II immunogenicity experiments, we followed the method by Kim et al.^37^ to classify pMHC-II with contradictory results as binding pairs. Moreover, we identified proteins with sequence similarity below 0.5 in the RCSB Protein Data Bank (RCSB PDB)^66^ and generated ten times of negative samples by randomly splitting peptides of the same length as the positive samples. The strategy aligns with the approaches used in IEPAPI^67^ and MHCflurry 2.0^68^. Subsequently, we randomly divided the data into training/validation and test sets at an 8:2 ratio. Consequently, the training/validation set comprised 71,584 data points, and the test set included 17,897 data points for our immunogenicity analysis.

#### Simulated multi-allelic dataset

Due to the absence of precise labels in the multi-allelic data, we constructed a simulated multi-allelic dataset using the single-allelic dataset, which has been divided into a 5-fold cross-validation set, to evaluate the capability of MIL module in detecting positive pMHC-II samples within bags. The process was as follows: We selected four out of the five folds as the training set. This data was then randomly shuffled and organized into bags, each containing ten samples. Subsequently, we randomly sampled negative instances to achieve a 1:3 ratio of positive to negative bags.

### ImmuScope architecture

#### Multi-allelic and single-allelic data representation

In this study, our model processed two predominant forms of mass spectrometry immunopeptidomics data: multi-and single-allelic data. Following the paradigm of multiple-instance learning, we treated each multi-allelic sample as a “bag” containing multiple instances, specifically pMHC-II pairs (**Supplementary Fig. 6c**). A positive bag suggests that the peptides are presented by at least one of the MHC molecules expressed in that sample. Conversely, a negative bag indicates that all pMHC-II pairs are negative instances. Similarly, for single-allelic data, we defined each pMHC-II sample as either a positive bag with a single positive instance or a negative bag with a single negative instance. This consistent representation of multi- and single-allelic data enabled our framework to simultaneously learn from both data types and make predictions, thereby facilitating its application across diverse immunopeptidomics datasets.

#### Attention-based MIL aggregator

In *branch a* of **Fig. 1a**, we employed an attention-based MIL pooling mechanism^69, 70^ to aggregate instance features within each bag. This mechanism not only enhances interpretability for predicting bag labels but also enables the identification and prioritization of the most critical instances crucial for the final prediction. Let z= ɛ (**x**; θ) represent the embedding of pMHC-II instance obtained from the backbone of ImmuScope ɛ parameterized by θ. **z***_k_* denotes the *k*-th instance in the bag ***z*** = {***z***_1_,…,**z***_K_*}. We implemented the following gated attention aggregator:

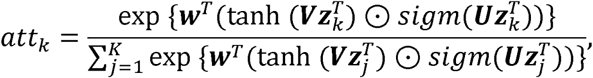

where ***w***, ***v*** and ***U*** denote model parameters, ⊙ represents an element-wise multiplication. The function *tanh*(·) refers to the hyperbolic tangent activation function and *sigm*(·) denotes the sigmoid non-linearity.

#### High-confidence positive pseudo-labels selection module

We introduced high-confidence positive pseudo-labels to improve the accuracy of antigen presentation prediction. We observed that the volume of positive samples in multi-allelic data was approximately twice that of single-allelic data, and the coverage of alleles was 2.2 times larger. This difference is particularly evident at the HLA-DP and HLA-DQ loci, where multi-allelic data substantially supplements the coverage gaps in single-allelic data. Notably, these weakly annotated positive multi-allelic samples contain multiple pMHC-II pairs, with at least one pair exhibiting positive signals. Such characteristics pose challenges for directly incorporating multi-allelic data into model training. To address this, we have developed a high-confidence positive pseudo-label selection module, which self-iteratively incorporates positive pseudo-labels from multi-allelic data to refine our predictive model (**Fig. 1f**).

The selection of high-confidence positive samples is conducted using the trained backbone of ImmuScope, with the specific training process detailed in the ***antigen presentation prediction*** section. Initially, multi-allelic data are input into the backbone of ImmuScope, which incorporates Monte-Carlo Dropout (MC Dropout)^71^ to assess variability and enhance reliability. Following this, an attention-based MIL aggregation module is used to obtain the uncertainty distribution of the multi-allelic samples, enabling the identification of high-confidence positive samples. Specifically, we iteratively select high-confidence samples from the multi-allelic data by controlling the confidence ratio (Top R%) associated with the attention scores. Samples that exhibit high confidence within the antigen presentation prediction branch are concurrently excluded from selection. The identified high-confidence positive samples are then integrated into the single-allelic data for model fine-tuning. Throughout this iterative process, we progressively adjust the confidence thresholds to incorporate a wider range of positive multi-allelic samples, thereby enhancing the model’s generalization capabilities. Finally, we identify the optimal ratio of positive pseudo-labeled samples by selecting the configuration that demonstrates the best performance on the validation set.

#### Positive-anchor triplet loss

MHC-II molecules exhibit significant diversity, exemplified by human MHC-II such as HLA-DR, HLA-DQ, and HLA-DP, which collectively display a remarkable total of 11,674 allelic variants according to the IPD-IMGT/HLA Database^72^. Additionally, the peptides themselves demonstrate notable variability in sequence and length. The peptide-binding groove of MHC-II molecule is highly specific for binding to amino acids in peptides^73^, determining which peptides can be bound and presented. Triplet loss^74^ enhances the model’s ability to perceive these subtle differences by minimizing the distance between similar samples (positive samples) and maximizing the distance between dissimilar samples (negative samples). This loss is particularly suitable for predicting pMHC-II binding affinity and antigen presentation, as it amplifies the learning efficacy for challenging-to-discriminate pMHC-II samples, thereby facilitating the discovery of more nuanced binding patterns between peptides and specific MHC-II molecules.

In the experimental setup, we calculated the triplet loss using only positive samples as anchors. This strategy enabled the model to more effectively distinguish crucial binding features within pMHC-II complexes. Moreover, the ratio of positive to negative samples in the antigen presentation dataset was set to 1:10. Using negative samples as anchors increased computational costs and might distract from the model’s primary goals by unnecessarily optimizing distances between negative samples. Such optimizations failed to enhance the model’s discriminative capabilities and reduced learning efficiency. To address these challenges and align with critical learning objectives, we have formulated the triplet loss for each mini-batch as follows:

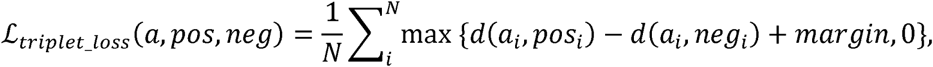

Where *d*(*x_i_*,*y_i_*) = |||*x_i_* - *y_i_*||*_P_*, we used Euclidean distance as the metric function, setting *p* = 2. In this context, *i* represents a mini batch, *N* is the batch size, and *a* exclusively denotes all positive samples used as anchors. *pos* and *neg* indicate the positive and negative samples within the mini-batch, respectively. The *margin* is a threshold defining the minimum distance that the negative sample must exceed beyond the positive sample from the anchor to avoid incurring a loss.

### ImmuScope Training process

#### ImmuScope backbone training process

The backbone of ImmuScope is a pre-trained model for other downstream tasks. Initially, we loaded the single- and multi-allelic data, and then we computed the positive-anchor triplet loss for the embeddings of pMHC-II instances, denoted as *L_triplet_loss_*. In *branch a*, the bag labels for single- and multi-allelic data were optimized using the binary cross entropy (BCE) loss function, represented as *L_MIL_MA_*, *L_MIL_SA_,* respectively. Concurrently, in *branch b*, the single-allelic data was optimized using the BCE loss *L_instance_SA_*. The composite loss function for the backbone is defined as:

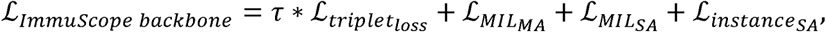

where *τ* represents the weighting factor for the triplet loss, setting *τ*= 0.1. Throughout the training process, the parameters of the ImmuScope backbone network were refined by synergistically combining individual instance learning, aggregated label optimization, and metric learning strategies. This integrative approach ensured a robust optimization of model parameters, effectively capturing both micro and macro-level data characteristics. The Adam optimizer with a learning rate of 1x 10^-3^ was used to train the backbone of ImmuScope for up to 20 epochs, with the final model being selected based on the best performance on the validation set.

#### Antigen Presentation Prediction

Based on the backbone of ImmuScope, we gradually introduced high-quality positive pseudo-labels from multi-allelic data to construct an antigen presentation prediction model. In each epoch, we first obtained the predicted antigen presentation probability on *branch b*, the attention score in the MIL aggregator, and the corresponding bag score through forward calculations. To ensure prediction stability and accurately gauge model uncertainty, we employed an architecture with MC Dropout to perform ten forward passes and analyzed both the mean and variance of these predictions. Initially, we selected the top 8% of samples with high attention weights and whose variances ranked in the top 80% (from lowest to highest). These thresholds (8% and 80%) were determined through preliminary experiments and an examination of the distribution of attention scores, ensuring that we focused on high-confidence, relatively low-variance samples. We also excluded samples with predicted antigen presentation probabilities exceeding 0.95 and those whose variances ranked in the top 40% (from lowest to highest), as they were already reliably identified by the model.

As the iterations progressed and the model’s internal representations became more stable, we gradually relaxed the threshold on attention weights from the top 8% to 12%. This step—commonly employed in self-training approaches—aims to broaden the scope of positive pseudo-labeled samples, thereby enriching the training dataset with more diverse pMHC-II binding candidates and further enhancing the model’s learning capacity. At the same time, we utilized this expanded dataset, SA-extend (EL), for incremental fine-tuning of the backbone model. Finally, we fine-tuned the ImmuScope backbone with the final SA-extend (EL) dataset over 10 additional epochs using the Adam optimizer (learning rate =3 x 10^-5^), producing the optimized ImmuScope-EL model for antigen presentation prediction.

#### CD4^+^ T cell epitope prediction

The antigen presentation process is the prerequisite for CD4^+^ T cell immune response. In line with the methodology of NetMHCIIpan-4.3, our CD4^+^ T cell epitope prediction model, ImmuScope, similarly incorporated both BA and EL data. Specifically, the antigen presentation model ImmuScope-EL was fine-tuned using binding affinity data, employing a learning rate of 2x 10^-5^, and leveraging the Adam optimizer to minimize the Mean Squared Error (MSE) loss over 20 epochs. To balance the influence of BA and EL data on CD4^+^ T cell epitope prediction, we set an 8:2 weighting ratio for binding affinity and single-allelic data in the validation set. This ratio was determined based on preliminary experiments and data correlation: while binding affinity data provide precise binding affinity information, single-allelic data capture actual antigen presentation events in *vivo*. The final validation metrics were calculated as follows:

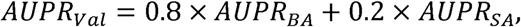

where *AUPR_BA_* and *AUPR_SA_* denote the AUPR values of the binding affinity and single-allelic subsets, respectively, within the validation set. Finally, we evaluated the performance of CD4^+^ T cell epitope prediction on the CD4^+^ epitope benchmark.

#### MHC-II epitope immunogenicity prediction

Immunogenicity is crucial as it determines the efficacy and safety of vaccines and therapies by triggering immune responses. We refined ImmuScope-EL further with immunogenicity data to develop the ImmuScope-IM model, tailored to immunogenicity prediction. The ImmuScope-IM model was optimized by an Adam optimizer with a learning rate of 1x 10^-3^ and Binary Cross-Entropy Loss (BCELoss), for a maximum of 20 epochs. For the application of the ImmuScope-IM model in SARS-CoV-2 epitope discovery and dynamic escape mechanism studies, we excluded the epitope binding data pertaining to SARS-CoV-2 from our initial immunogenicity dataset to construct a dedicated SARS-CoV-2 immunogenicity benchmark dataset, ensuring unbiased benchmarking. This benchmark dataset was then used to train the ImmuScope-IM model for assessing SARS-CoV-2 epitope.

In this study, all deep learning models were developed utilizing PyTorch version 1.12.1 and were trained utilizing an NVIDIA GeForce RTX 4090 GPU. Comprehensive details on the algorithm and model hyperparameters are enumerated in **Supplementary Table 5** and **6**, respectively.

### Analysis of motif deconvolution

We employed the trained ImmuScope-EL model to perform motif deconvolution and obtain the binding peptide sequence set for different MHC-II allomorphs. Specifically, a specific subset of multi-allelic data was input into ImmuScope-EL, and then the weight of attention-based MIL aggregator and the antigen presentation probability were obtained from *branch a* and *branch b* respectively. To ensure high-quality deconvolution, we selected the antigen presentation peptides with an antigen presentation probability greater than 0.8 and an attention weight exceeding the reciprocal of the number of MHC-II categories in the bag. We then employed Seq2Logo to visualize the motif logo of different MHC-II allomorphs based on the sequences of selected peptides.

### Quantification of MHC-II binding specificity

We first calculated the antigen presentation score by inputting 100,000 random human peptide sequences and the alleles to be assessed into ImmuScope-EL. Then, the samples with the top 1% of the predicted scores were selected for cluster analysis using GibbsCluster^75^, and the optimal number of clusters, i.e., binding specificity, was determined based on the lowest average KLD. Finally, we evaluated the MHC binding specificity quantified by ImmuScope-EL by comparing the KLD with the PSFM matrix based on the peptidomics data. The prediction results of NetMHCIIpan were obtained by predicting the top 1% of random human peptides using the NetMHCIIpan-4.3 software package, while MixMHC2pred was obtained by predicting using the MixMHC2pred-2.0 webserver.

### Measuring the similarity of MHC binding motifs

To evaluate the similarity between sequence motifs generated by various algorithms and those obtained from peptidomics data, we first represented each set of peptide-binding cores with PSFMs. Each PSFM was then converted into a single vector by concatenating the frequency values at its nine positions, with each position containing 20 values corresponding to the 20 standard amino acids. Finally, we calculated the symmetric Kullback-Leibler Divergence^18^ for any two PSFM vectors *a* and *b* using the following formula:

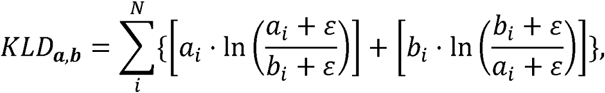

where *ɛ* is employed as an exceedingly small positive number, typically set at 1x 10^-10^, to prevent division by zero.

### Calculation of binding core alignment scores for epitopes

In our analysis of the melanoma neoepitope and SARS-CoV-2 epitope binding cores, we employed the ImmuScope-EL model to analyze the binding cores of various epitopes and to examine changes upon mutations. Initially, we used ImmuScope-EL to predict 100,000 random human peptides and selected the top 1% based on the highest binding scores to create a position-specific scoring matrix (PSSM) for specific alleles (like HLA-DRB1*01:01 in SARS-CoV-2 epitope analysis). Subsequently, we calculated the matching degree for each 9-length window of the candidate peptides against the PSSM. The alignment score for each window was then computed to assess how well the subsequence of the window matched the binding modality.

### Statistical analyses

Error bars depicted in the bar plots indicate 95% CIs, unless specified otherwise. Performance benchmarks such as AUC and AUPR were computed using the scikit-learn Python package, version 1.3.0. UMAP analysis was conducted with the umap-learn Python package, version 0.5.3. The predicted binding peptide ligands were further clustered using the GibbsCluster tool, version 2.0. Sequence motifs were generated and visualized using the Seq2Logo tool, version 2.0. Additionally, 3D structures of pMHC-II complexes were visualized using PyMOL, version 2.5.7.

## Data availability

The MHC-II antigen presentation data and CD4^+^ epitope benchmark are sourced from NetMHCIIpan-4.3^18^ (https://services.healthtech.dtu.dk/services/NetMHCIIpan-4.3/). Immunogenicity datasets are derived from IEDB^63^ (https://www.iedb.org/). The motifs of MHC-II alleles are obtained from the MHC Motif Atlas Database^47^ (http://mhcmotifatlas.org/) and NetMHCIIpan-4.3. Melanoma neoepitope immunogenicity data are derived from original studies^52^ (https://doi.org/10.1038/s41586-022-04682-5). Data on SARS-CoV-2-derived CD4^+^ T cell epitopes, the structure of HLA class II-presented SARS-CoV-2 epitopes, and other metadata regarding SARS-CoV-2 escape from CD4^+^ T cells are sourced from the original study^53^ (https://doi.org/10.1016/j.celrep.2023.112827). HLA class II sequences are obtained from IPD-IMGT/HLA^76^ (https://www.ebi.ac.uk/ipd/imgt/hla/). Data pertaining to the sequence evolution of SARS-CoV-2 variants are derived from ViralZone^77^ (https://viralzone.expasy.org/). Source data are provided with this paper.

## Acknowledgements

This work was supported by the National Natural Science Foundation of China (grant nos. 62372234, 62072243) and the Postgraduate Research & Practice Innovation Program of Jiangsu Province (KYCX23_0490). Funding was also provided by the National Health and Medical Research Council of Australia (grant nos. APP1127948, APP1144652, and APP2036864 to J.S.), and by the Major and Seed Inter-Disciplinary Research projects from Monash University (J.S.).

## Author contributions

L.-C.S., Y.Z., D.-J.Y., and J.S. conceived the concept. L.-C.S. and Z.W. designed the computational architecture. L.-C.S. conducted the computational experiments and developed the downstream tasks. L.-C.S. and Y.Z. performed the bioinformatics analyses. L.-C.S. prepared the figures and wrote the manuscript. D.R.L., Y.L., J.T., and J.R. provided guidance on data analysis. D.-J.Y. and J.S. supervised the project. All authors contributed ideas to the work and participated in editing and revising the manuscript.

## Extended data figure legends

**Extended Data Fig. 1.**
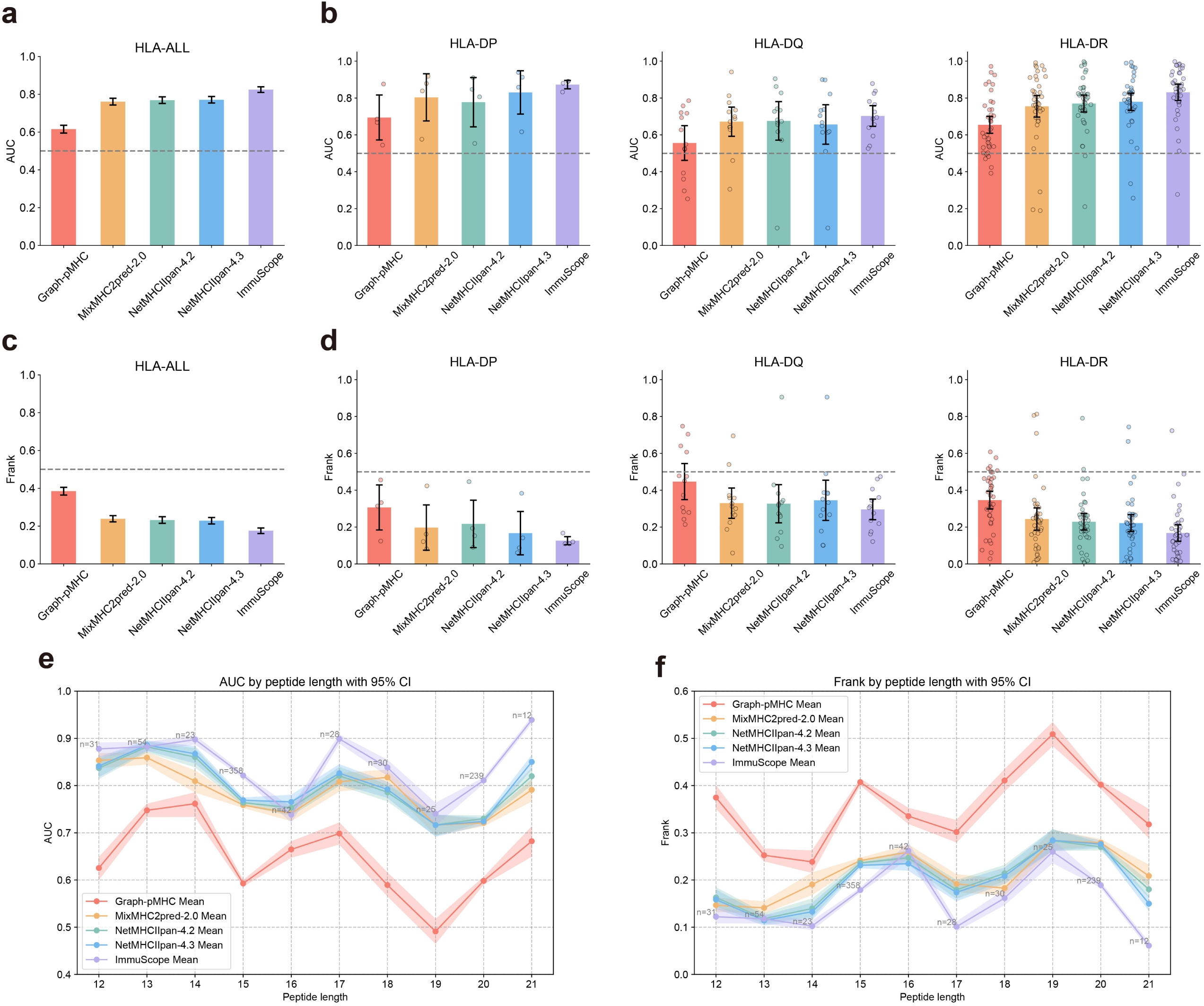
ImmuScope improves CD4^+^ T cell epitope prediction. **a**, Bar plot of AUCs on the CD4^+^ epitope benchmark. **b**, Bar plots of AUCs at different HLA loci (HLA-DR, HLA-DP and HLA-DQ), the points represent the average AUC of the corresponding MHC molecule. **c**, Bar plot of Frank values on the CD4^+^ epitope benchmark. **d**, Bar plots of Frank values at different HLA loci. In **a**-**d**, the bars represent the mean AUCs or Frank values by 1,000 bootstrap iterations, the error bars indicate the 95% CIs, and each data point represents the performance of the corresponding HLA loci. **e,f,** Comparison of AUC (e) and Frank (f) performance between ImmuScope and other methods on the CD4^+^ epitope benchmark across different peptide lengths.

**Extended Data Fig. 2.**
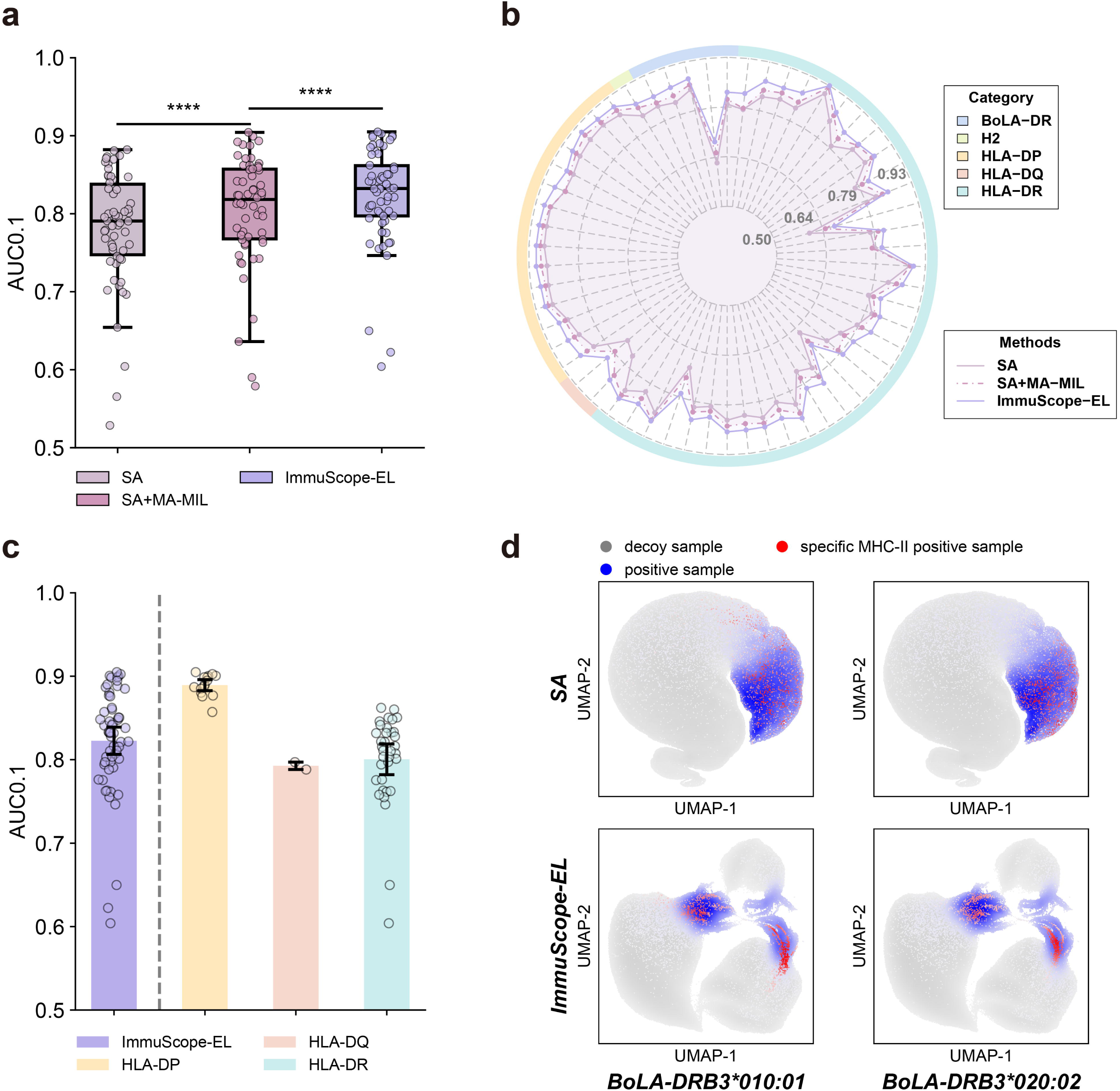
Ablation experiments on key modules of ImmuScope-EL. **a**, AUC0.1 performance of SA, SA+MA-MIL, and ImmuScope-EL. Box center line, median; box limits, upper and lower quartiles; whiskers, 1.5×interquartile range; ****P < 0.0001. Each data point represents the performance of the corresponding MHC allele. **b**, AUPR performance of SA, SA+MA-MIL, and ImmuScope-EL, calculated per MHC-II allele, only including alleles with at least 25 positive samples and a minimum of 30 total samples. **c**, AUC0.1 performance of the ImmuScope-EL model for different MHC alleles. The bars represent the mean performance values by 1,000 bootstrap iterations, and the error bars indicate the 95% CIs. Each data point represents the performance of the corresponding MHC allele. **d**, UMAP visualization of instance embeddings from the SA and ImmuScope-EL models on the BoLA-DRB3 subset.

**Extended Data Fig. 3.**
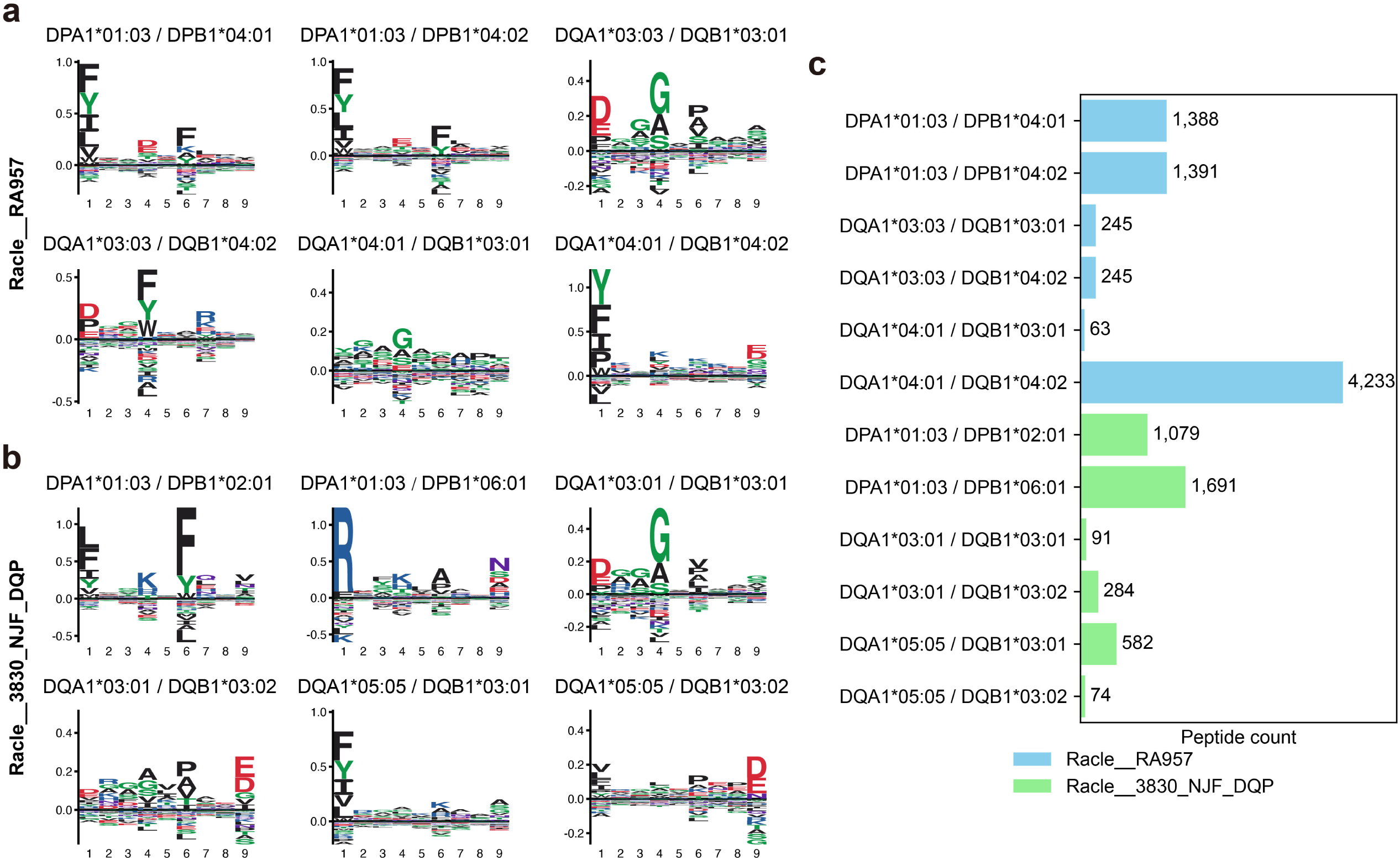
Deconvolution of ImmuScope on the multi-allelic data. **a**, **b**, Motif deconvolution logo on Racle RA957 (**a**) and Racle 3830_NJF_DQP (**b**) datasets. **c**, Motif deconvolution peptide count on the heterozygous subsets.

**Extended Data Fig. 4.**
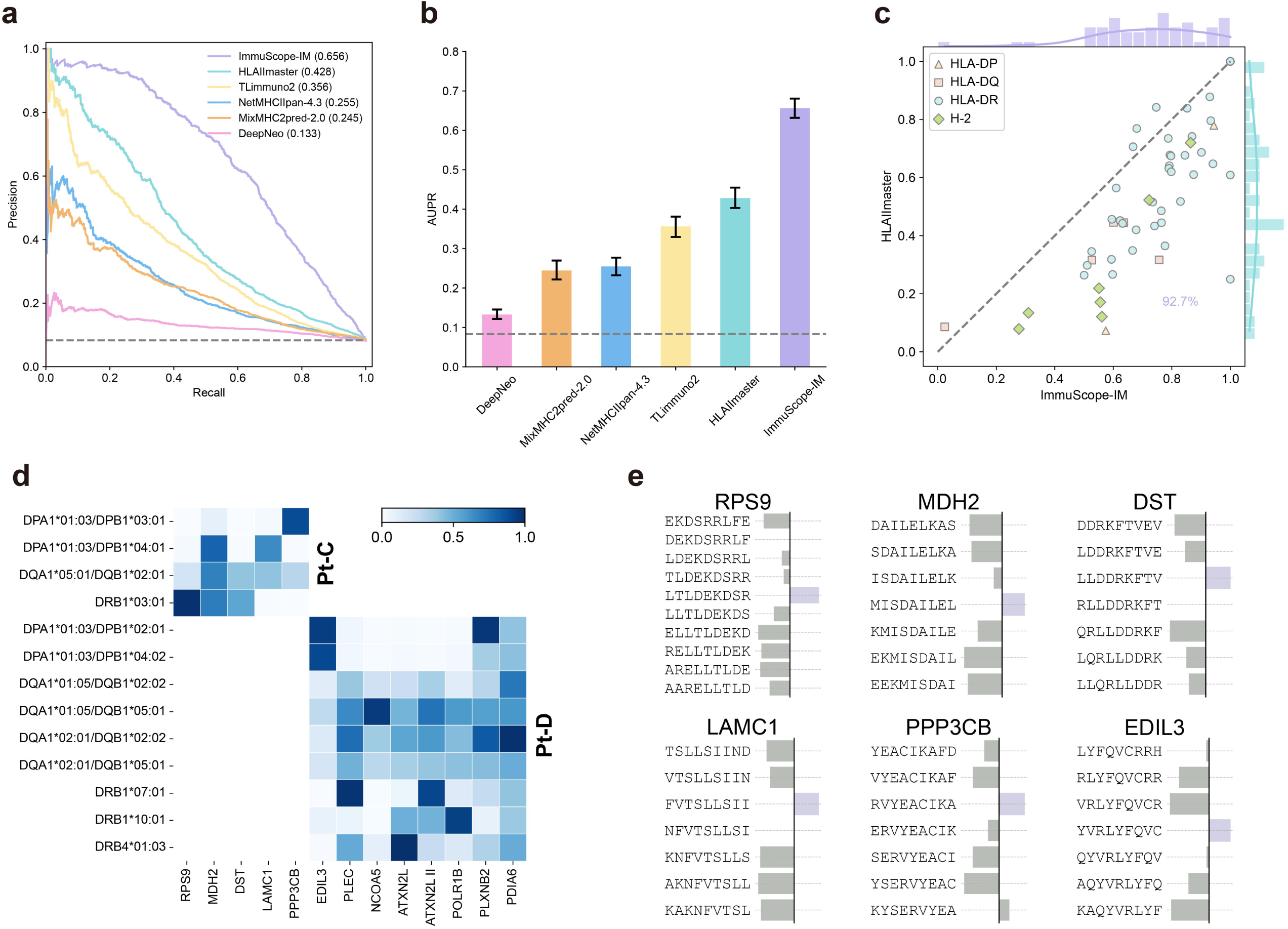
Comparison of predictive performance for immunogenicity benchmarking. a, **b**, Precision-recall curves (**a**) and bar plots of AUPRs (**b**) of the candidate methods on the immunogenicity benchmark. The error bars indicate the 95% confidence intervals by 1000 bootstrap iterations. The gray dashed lines denote the results of random predictions. **c**, AUPR performance comparison between HLAIImaster and ImmuScope-IM on different MHC alleles. **d**, Predictive analysis of HLA restriction for melanoma neoantigens. The x-axis corresponds to different neoantigens, and the y-axis represents different HLA alleles. Each position represents the antigen presentation probability of the corresponding antigen and allele predicted by ImmuScope-EL. The upper left matrix corresponds to Pt-C, and the lower right matrix corresponds to Pt-D. The color depth corresponds to the antigen presentation probability. **e**, Predicted cores and alignment scores for melanoma neoantigen-HLA binding with ImmuScope-EL. Purple represents high alignment scores.

**Extended Data Fig. 5.**
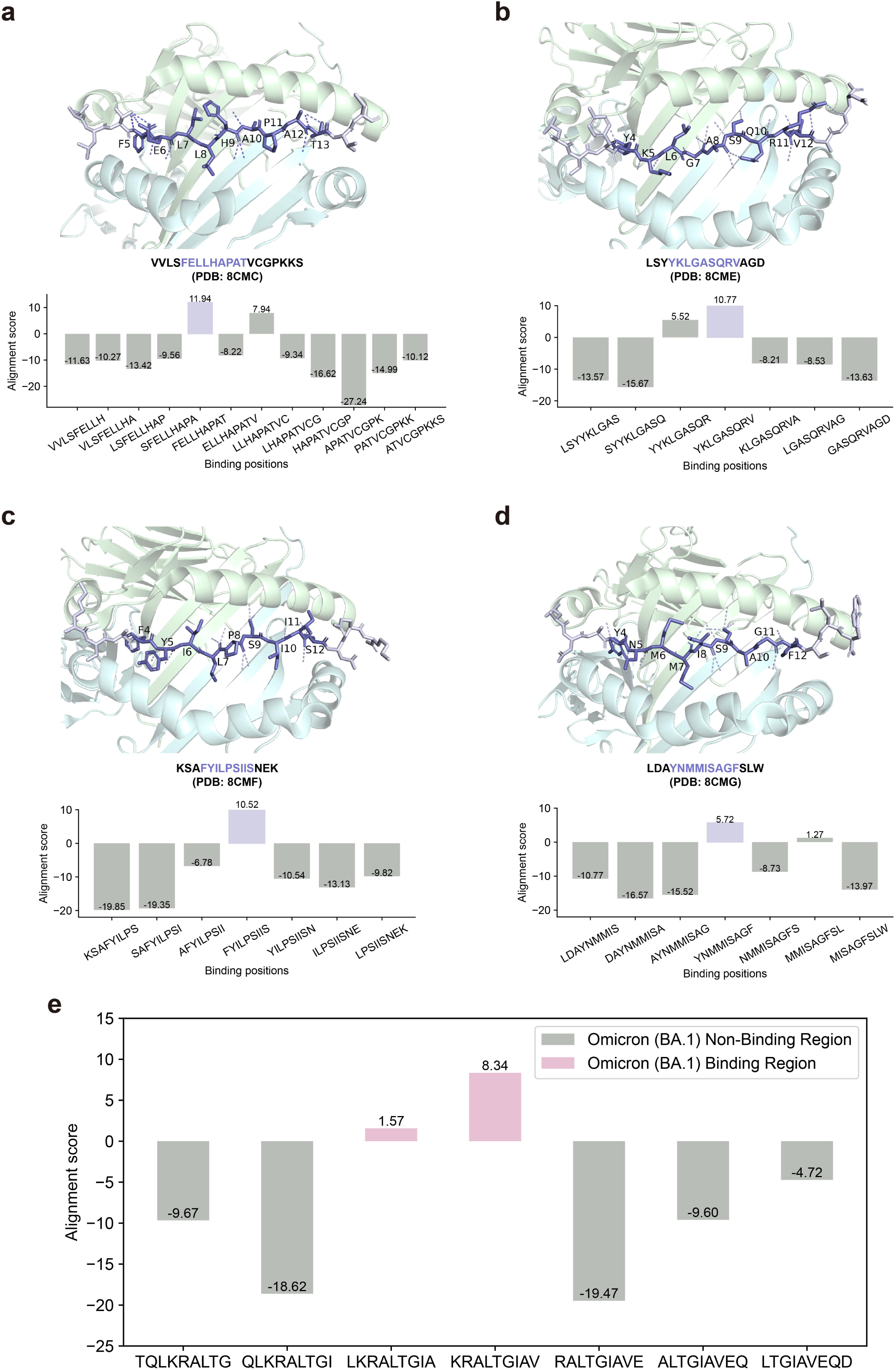
Predicted binding position by ImmuScope-EL in SARS-CoV-2 spike protein epitopes. **a**, **b**, **c**, **d**, binding positions and alignment scores on HLA-DR1-S_511-530_ (PDB: 8CMC, **a**), HLA-DR1-M_176-190_ (PDB: 8CME, **b**), HLA-DR1-nsp3_1350-1364_ (PDB: 8CMF, **c**) and HLA-DR1-nsp14_6420-6434_ (PDB: 8CMG, **d**), respectively. **e**. Alignment scores for various binding positions of HLA-DR1-S_761-775_^Omicron^ ^(BA.1)^ as predicted by ImmuScope-EL excluding position P5.

## Supplementary figure and table legends

**Supplementary Fig. 1** Detailed model architecture of ImmuScope backbone.

**Supplementary Fig. 2** Ablation experiments on key modules of ImmuScope-EL.

**Supplementary Fig. 3** Motifs from the MHC Motif Atlas Database and NetMHCIIpan-4.3.

**Supplementary Fig. 4** Binding motifs predicted by ImmuScope-EL.

**Supplementary Fig. 5** Predicted cores and alignment scores for melanoma neoantigen-HLA binding with ImmuScope-EL.

**Supplementary Fig. 6** Overview of MHC-II antigen presentation datasets and data organization.

**Supplementary Table 1** Basic characteristics of discovery cohort.

**Supplementary Table 2** HLA alleles of discovery cohort.

**Supplementary Table 3** Statistics of AlphaFold3-predicted pMHC complex structures.

**Supplementary Table 4** Statistical overview of the datasets used for model training in this study.

**Supplementary Table 5** Statistical summary of ImmuScope algorithm hyperparameters.

**Supplementary Table 6** Statistical summary of ImmuScope model hyperparameters.

